# SPATNIC enables H&E-grounded cancer-cell identification across spatial and single-cell transcriptomic modalities in colorectal cancer

**DOI:** 10.64898/2026.10.21.760612

**Authors:** Shunsuke A. Sakai, Ryosuke Nomura, Masaki Ohira, Kyoko Kurihara, SungGi Chi, Yuki Matsubara, Yukiko Morinaga, Takeshi Kato, Naoko Takeda-Miyata, Motohiro Kojima, Shingo Sakashita, Hiromichi Ebi, Rui Yamaguchi, Eiji Oki, Takao Fujisawa, Hideaki Bando, Ayako Suzuki, Yutaka Suzuki, Katsuya Tsuchihara, Shun-Ichiro Kageyama, Riu Yamashita

## Abstract

Tumors contain diverse non-cancer cells that constitute the tumor microenvironment; thus, identifying which cells are cancerous is a foundational and critical first step in cancer research. However, to our knowledge, no transcriptome-based classifier has been trained using pathologist-defined annotations from matched H&E images at single-cell resolution. Moreover, few tools can be shared across diverse platforms. Here, we present SPATNIC, a variational autoencoder that classifies individual cells as cancer or normal, trained on Xenium cells labeled by a pathologist review of matched H&E images. It matches or exceeds the performance of existing tools in primary colorectal tumors and liver and lung metastases. SPATNIC generalizes zero-shot across Visium, Visium HD, CosMx, Xenium, MERSCOPE, and single-cell RNA sequencing. In public single-cell atlases, it identified a rare *POU2F3*⁺ tuft-like cancer stem cell population missed by existing annotations and associated with worse survival. In rectal cancer profiled before and after chemoradiotherapy, it separated cancer from normal epithelium in tissue, thus enabling per-patient screening of antibody–drug conjugate (ADC) targets.

## Introduction

Distinguishing cancer cells from their normal counterparts is fundamental in cancer research. Current single-cell and spatial transcriptomics profile gene expression at cellular or near-cellular resolution, with spatial platforms preserving tissue architecture, thereby enabling the study of cancer cells within their native microenvironment^1^. Thus, reliable classification of individual cells as cancer or normal is essential. This requirement has motivated the development of a series of computational classifiers; however, the task remains incomplete^2^.

The current approaches face three limitations. The first concerns the labels. Early methods identified cancer cells by inferring copy-number variation from gene expression—inferCNV^3^, copyKAT^4^, CaSpER^5^, SCEVAN^6^, and Numbat^7^—which generally depends on broad transcriptome coverage and frequently on a normal reference, which limits their transferability to targeted panels. More recent machine-learning classifiers have moved beyond copy-number inference: CancerFinder^8^ applies domain generalization to a feed-forward network to facilitate transfer across datasets, scMalignantFinder^9^ and ikarus^10^ classify or score cells using logistic regression with curated cancer signatures or differentially expressed genes, and scATOMIC^11^ classifies cells hierarchically across many cancer types. Despite these advances, these methods have been developed and evaluated predominantly using single-cell RNA sequencing (scRNA-seq) data, with malignant cell labels typically based on prior annotations, inferred genomic alterations, or expression signatures, rather than direct histopathological assessment of matched tissue.

The second limitation is organ dependence. Although the classification is binary—cancer or normal— both malignant and non-malignant transcriptional states vary across anatomical sites. For example, the non-malignant epithelial compartment comprises colonic mucosa at the primary colorectal site, hepatocytes and cholangiocytes in the liver, and alveolar and bronchial epithelial cells in the lungs. Few classifiers explicitly account for the anatomical site, and we reasoned that not accounting for these tissue-specific transcriptional contexts may reduce accuracy when a single model is applied across primary tumors and metastatic sites. The third limitation is platform dependence. Single-cell and spatial transcriptomic technologies currently span various platforms—Visium^12^, Visium HD^13^, CosMx^14^, Xenium^15^, MERSCOPE^16^, and scRNA-seq^17^, which differ in gene coverage, detection sensitivity, and spatial resolution, and laboratories adopt them unevenly^18^. To our knowledge, no cancer cell classifier has been systematically validated across this range of transcriptomic platforms, potentially rendering method selection dependent on the data type.

Here, we introduce SPATNIC (Spatial Pathology-based Typist for Normal/Cancer), a variational autoencoder that classifies individual cells as cancer or normal based on gene expression, trained on cells annotated through pathologist review of matched H&E-stained tissue rather than on expression-derived labels. Built as organ-specific models for primary colorectal tumors and their liver and lung metastases, SPATNIC accurately identifies cancer cells and matches or exceeds existing classifiers, while achieving greater computational efficiency. We show that, within the metastatic site, a single SPATNIC model generalizes without platform-specific retraining across Visium, Visium HD, CosMx, Xenium, MERSCOPE, and scRNA-seq, and remains robust to substantial reductions in gene-panel content. We then applied SPATNIC in two settings: public scRNA-seq atlases^19,20,21^, where discrepancies between SPATNIC predictions and existing annotations identified a rare cancer cell subset associated with worse survival, and to rectal cancer samples profiled before and after chemoradiotherapy, where SPATNIC distinguished cancer cells from normal epithelium cells within the tissue and identified candidate patient-specific targets for ADC targets.

## Results

### SPATNIC accurately distinguishes cancer from normal cells across primary and metastatic colorectal cancer

SPATNIC uses the gene expression profile of a single cell as input and returns the probability of the cell being cancer rather than normal (Fig. 1a). It encodes each cell’s expression profile into a latent representation (*z* in Fig. 1a) using a Student-*t* observation model, whose heavy tails reduce sensitivity to extreme and highly variable counts in sparse single-cell and spatial transcriptomic data. The classification head then maps this representation to the probability of cancer. Because the model uses only gene expression as input, it requires neither copy-number inference nor a matched normal reference.

**Figure 1.**
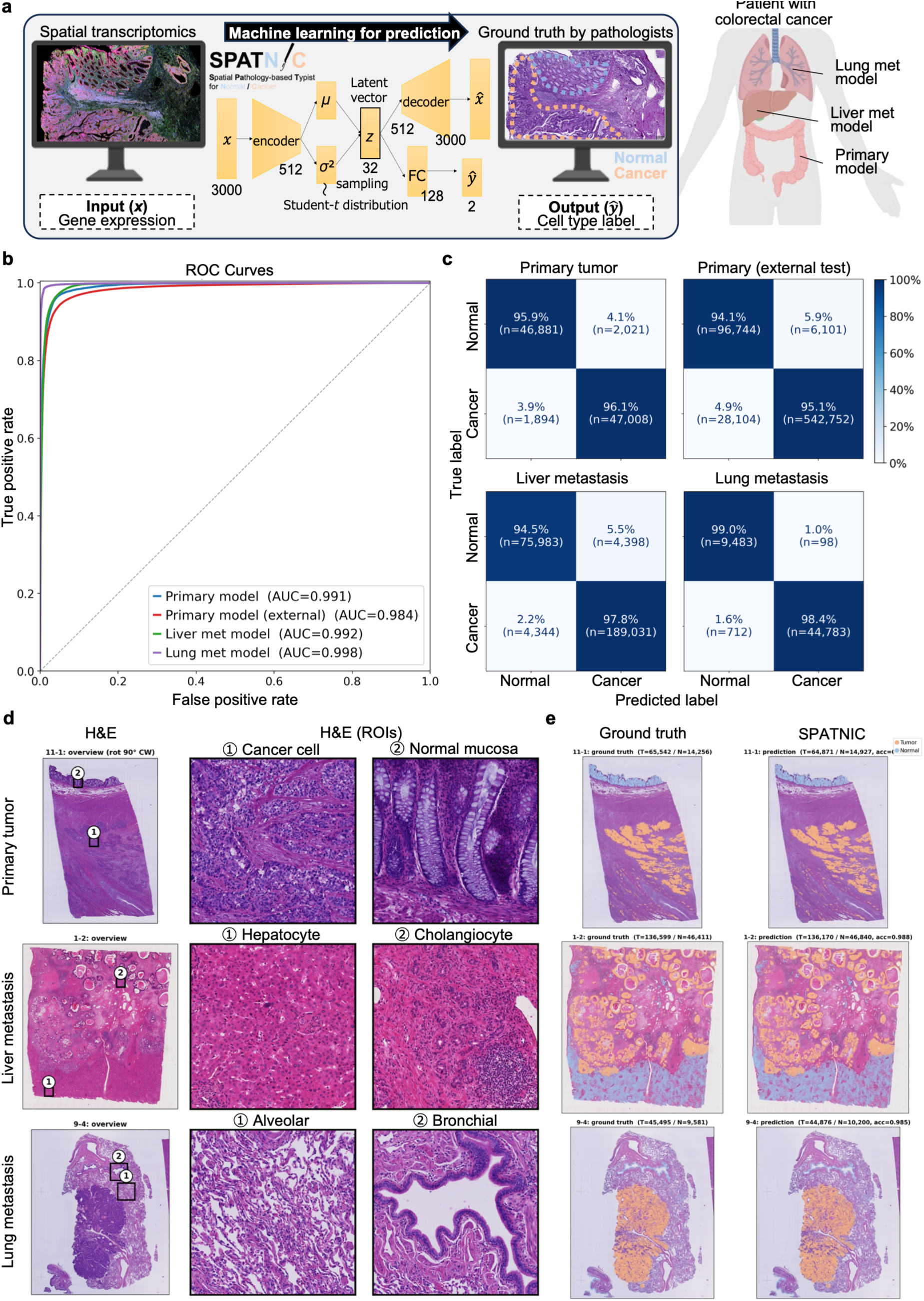
SPATNIC accurately classifies cancer and normal cells across primary and metastatic colorectal cancer. (a) Schematic of the SPATNIC framework. Spatial transcriptomic gene-expression profiles (input, x) from colorectal cancer (CRC) primary tumors and matched liver and lung metastases are used to predict a per-cell cancer/normal label (output, ŷ), benchmarked against pathologist ground-truth annotation. FC: fully connected layer. (b) Receiver operating characteristic (ROC) curves for the primary model (internal and external test sets) and the liver- and lung-metastasis models. Area under the curve (AUC) is indicated for each. (c) Confusion matrices for the primary tumor, primary external test, liver metastasis, and lung metastasis cohorts. Values indicate row-normalized percentages with absolute cell counts (n) in parentheses; color encodes percentage. (d) Representative H&E sections (left) with magnified regions of interest (ROIs) highlighting cancer cells and adjacent normal compartments (normal mucosa, hepatocytes/cholangiocytes, alveolar/bronchial epithelium) in primary tumor, liver metastasis, and lung metastasis. (e) Spatial overlays of pathologist ground truth and SPATNIC predictions for the sections in (d); cancer is shown in orange and normal in blue. Per-section cell counts (T, N) and accuracy are indicated.

We assembled a colorectal cancer (CRC) resource comprising rectal cancer Xenium samples with pathologist-annotated cancer/normal labels based on matched H&E staining, including a primary tumor cohort of 52 samples^22^ together with independent liver metastasis, lung metastasis, and external primary validation cohorts (Table 1). Three organ-specific models were trained—one each for primary tumors, liver metastases, and lung metastases—with class-balanced sampling during training (see the Methods section). Independent test sets were reserved for each anatomical site, and the external primary cohort was used exclusively to validate the primary model.

**Table 1.** Datasets and sample composition.

| Dataset | Training |  |  |  | Test |  |  |  |
| --- | --- | --- | --- | --- | --- | --- | --- | --- |
|  | Samples | Cells | Cancer | Normal | Samples | Cells | Cancer | Normal |
| Primary | 52 | 391,214 | 195,607 | 195,607 | 52 | 97,804 | 48,902 | 48,902 |
| Primary external † | 0 | 0 | 0 | 0 | 7 | 673,701 | 570,856 | 102,845 |
| Liver met ‡ | 12 | 200,000 | 100,000 | 100,000 | 3 | 273,756 | 193,375 | 80,381 |
| Lung met ‡ | 4 | 151,998 | 75,999 | 75,999 | 1 | 55,076 | 45,495 | 9,581 |

Numbers of samples and cells used for training and testing in each dataset, with the tumor/normal breakdown of cells. † External test set only; never used for training. ‡ Training data were used to fine-tune the primary model; test samples were held out at the sample level.

SPATNIC classified cancer and normal cells with high accuracy across all cohorts. On pooled cells, it achieved an AUROC of 0.991 and an AUPRC of 0.990 on the primary tumor test set, with comparable performance in the external primary cohort (AUROC 0.984, AUPRC 0.997), liver metastases (AUROC 0.992), and lung metastases (AUROC 0.998) (Fig. 1b, Table 2). The confusion matrices showed high sensitivity and specificity across cohorts, with no systematic bias toward either class (Fig. 1c, Table 2). Per-sample analysis showed a similar performance, while also revealing intersample variability across clinical specimens (Table S1).

**Table 2.** Pooled performance summary.

| Dataset | n cells | AUROC | AUPRC | MCC | F1 | Sensitivity | Specificity | Balanced acc. |
| --- | --- | --- | --- | --- | --- | --- | --- | --- |
| Primary | 97,804 | 0.991<br>[0.991–0.992] | 0.990<br>[0.990–0.991] | 0.920<br>[0.918–0.923] | 0.960<br>[0.959–0.961] | 0.961<br>[0.959–0.963] | 0.959<br>[0.957–0.961] | 0.960<br>[0.959–0.961] |
| Primary external | 673,701 | 0.984<br>[0.984–0.984] | 0.997<br>[0.997–0.997] | 0.825<br>[0.824–0.827] | 0.969<br>[0.969–0.970] | 0.951<br>[0.950–0.951] | 0.941<br>[0.940–0.942] | 0.946<br>[0.945–0.947] |
| Liver met | 273,756 | 0.992<br>[0.992–0.992] | 0.996<br>[0.995–0.996] | 0.923<br>[0.921–0.925] | 0.977<br>[0.977–0.978] | 0.978<br>[0.977–0.978] | 0.945<br>[0.944–0.947] | 0.961<br>[0.961–0.962] |
| Lung met | 55,076 | 0.998<br>[0.997–0.998] | 0.999<br>[0.999–1.000] | 0.930<br>[0.926–0.934] | 0.987<br>[0.986–0.988] | 0.977<br>[0.975–0.978] | 0.988<br>[0.986–0.990] | 0.982<br>[0.981–0.984] |

Values are point estimates with 95% bootstrap confidence intervals in brackets, computed over all pooled cells. n cells, number of pooled cells.

Because the composition of the non-malignant compartment differs substantially across anatomical sites, we next examined SPATNIC’s performance site by site in the tissue. On H&E, malignant glands were observed alongside normal mucosa in primary tumors, hepatocytes and cholangiocytes in liver metastases, and alveolar and bronchial epithelium in lung metastases (Fig. 1d). Despite this heterogeneity, spatial overlays of SPATNIC predictions onto the same tissue sections showed close concordance with the pathologist annotations in every organ (Fig. 1e), confirming the accurate classification of cancer and normal cells across primary and metastatic sites. To assess whether these organ-specific models could be applied beyond their training sites, we cross-applied each model to all the test cohorts (Tables S2 and S3). Each model performed the best within its corresponding anatomical site, producing diagonal dominance across the model × cohort matrix. These results support the use of organ-specific models, while also demonstrating partial cross-site transferability.

### SPATNIC learns interpretable features and supports confidence-aware classification

Next, we examined which genes contributed most strongly to the classification and found that the most discriminative features were biologically coherent (Fig. S1). The top contributors to the cancer class included glycolytic and hypoxia-associated genes (*SLC2A1*/GLUT1, *ALDOA*, *PGK1*, *VEGFA*)^23^, the chromatin regulator *HMGA1*^24^, the Wnt target *AXIN2*^25^, and *SLC40A1*, *FAT1*, and *GDF15*. Conversely, the top contributors to the normal class included canonical markers of intestinal identity and differentiation, including the master regulator *CDX1*^26^, goblet cell-associated genes such as *MUC4*, *MUC5B*, and *FCGBP*, and differentiation-associated factors *KLF4*^27^, *HOXB13*, and *CA12*. Representative markers showed concordant protein expression patterns according to immunohistochemistry (Fig. S1). This concordance indicates that SPATNIC relies on biologically interpretable features for classification.

SPATNIC also offers an optional strict-confidence mode that abstains on low-confidence cells. Relative to the default 0.5 threshold, this mode reduced both the false-negative (FN) and false-positive (FP) rates among the retained cells at the cost of classifying fewer cells (Fig. S2a, b). The abstained cells were enriched near the histological tumor–normal boundaries (Fig. S2c). Unless otherwise noted, all subsequent analyses used the default mode in which every cell was assigned a classification.

### SPATNIC outperforms existing classifiers in both classification performance and computational efficiency

We benchmarked SPATNIC against three published cancer cell classifiers—CancerFinder^8^, scMalignantFinder^9^, and ikarus^10^—each evaluated with its default pretrained weights and after retraining on our CRC Xenium data (Fig. 2a). In their default configurations, none of the three maintained useful classification Xenium data; across cohorts, their balanced accuracy was close to chance level, and ikarus assigned every cell to the normal class, yielding a Matthews correlation coefficient (MCC) of zero in the primary, external, and liver cohorts (Fig. 2a, Table S4). These tools were trained on scRNA-seq— dissociated cells profiled across the whole transcriptome, whereas Xenium profiled a targeted panel of 5,000 genes in intact tissue. The differences in gene coverage and measurement modalities may explain their limited transferability. SPATNIC, trained natively on Xenium 5K data, significantly outperformed all three baseline methods in their default configurations in per-sample AUC-PR across all four cohorts (P < 0.05, paired Wilcoxon signed-rank test; Fig. 2a, Table S5). Retraining the baselines on our data narrowed but did not close the gap. CancerFinder and scMalignantFinder achieved a performance comparable to SPATNIC once retrained on the primary cohort; for example, the retrained CancerFinder achieved an MCC of 0.909 versus SPATNIC’s 0.922, but neither exceeded it on any cohort (Table S4). Therefore, under the benchmark conditions used in this study, SPATNIC matched or exceeded the performance of the retrained baseline methods. Within SPATNIC itself, architecture ablation supported the choice of likelihood: replacing Student-*t* with a Gaussian one reduced tumor sensitivity in the external primary cohort (Fig. S3, Tables S6, S7). The spatial visualization of the predictions further illustrates these differences. Overlaid on tissue, SPATNIC closely recapitulated the pathologist annotations faithfully across primary tumors and metastases, whereas the default baselines diverged in characteristic ways: CancerFinder and scMalignantFinder overcalled cancer (high sensitivity, low specificity), whereas ikarus under-called almost throughout (Fig. S4).

**Figure 2.**
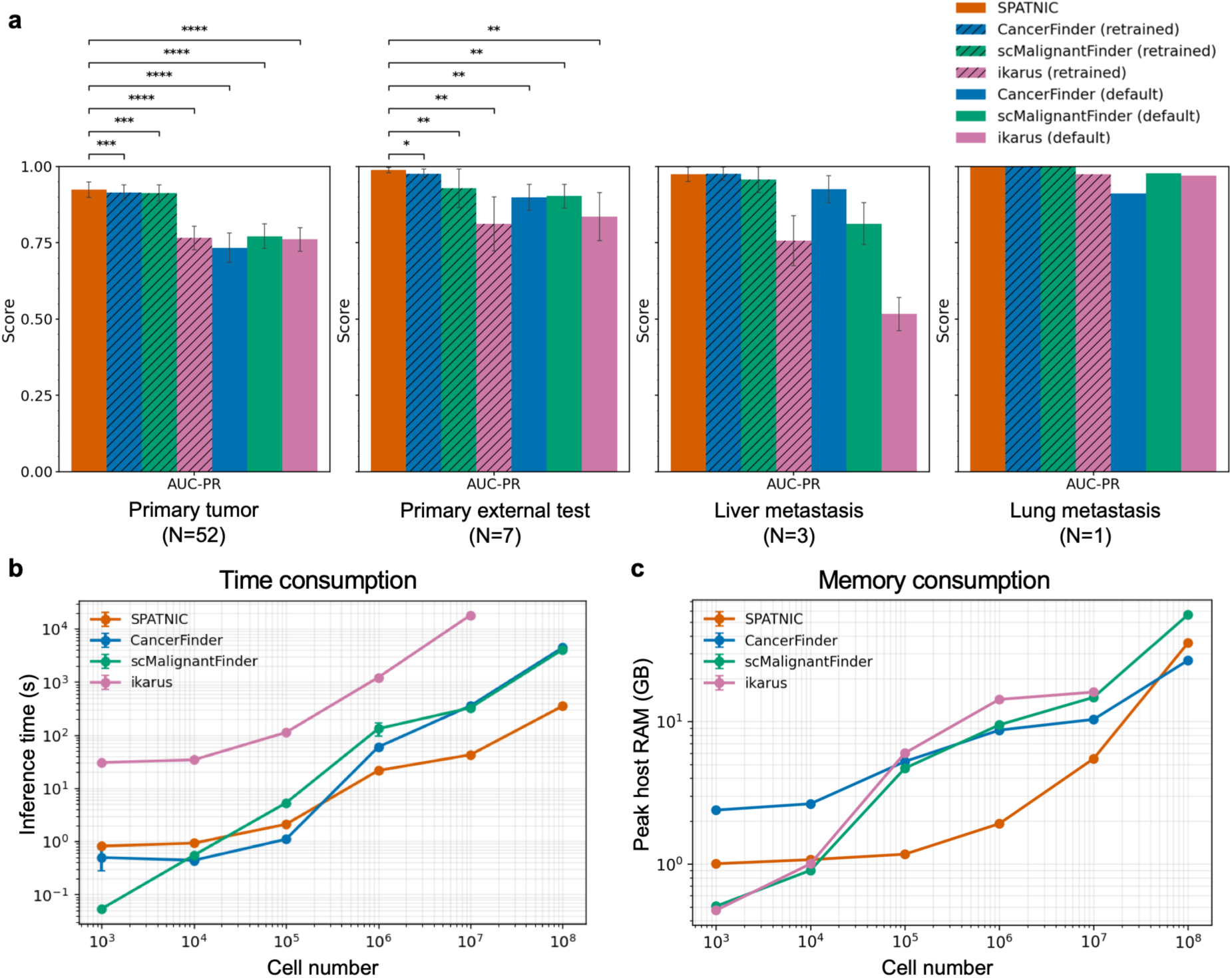
SPATNIC outperforms existing classifiers in accuracy and computational efficiency on Xenium data. (a) AUC-PR of SPATNIC versus CancerFinder, scMalignantFinder, and ikarus (each in retrained and default configurations) across primary tumor (N = 52), primary external test (N = 7), liver metastasis (N = 3), and lung metastasis (N = 1) cohorts. Bars show mean ± s.d.; asterisks denote significance versus SPATNIC (paired test): *P < 0.05, **P < 0.01, ***P < 0.001, ****P < 0.0001). (b) Inference time and (c) peak host RAM as a function of cell number for SPATNIC and benchmark methods, plotted on log– log axes.

In addition to its predictive performance, SPATNIC was substantially more efficient. Across increasing cell numbers, the inference time and peak memory footprint scaled favorably relative to the baselines, remaining tractable in scaling experiments using synthetic matrices of up to 10^8^ cells, whereas the competing methods exceeded the practical runtime or memory limits (Fig. 2b, c). Together, these results support SPATNIC as a more accurate and scalable method than the benchmarked methods for cancer cell identification in spatial transcriptomics.

### SPATNIC generalizes across spatial and single-cell platforms without platform-specific retraining

Spatial- and single-cell studies use platforms that differ widely in terms of gene coverage, measurement chemistry, and spatial resolution. Therefore, we assembled publicly available CRC datasets spanning Visium^12,28,29,30,31,32,33^, Visium HD^13^, CosMx^14^, Xenium V1^15^, MERSCOPE^16^, and scRNA-seq^17,19,20,21,34,35^, covering primary tumors and their liver and lung metastases, and applied SPATNIC without retraining or platform-specific fine-tuning (Fig. 3a). For each dataset, we categorized the available reference labels into one of three tiers: pathologist annotation of matched histology (histology-based reference); other published labels, such as author cell-type annotations (cell-type annotation); or our own manual annotation (self-annotated), where no reference label was available. The primary datasets were scored using the primary SPATNIC model, whereas the metastatic datasets were scored using the corresponding liver or lung metastasis models.

**Figure 3.**
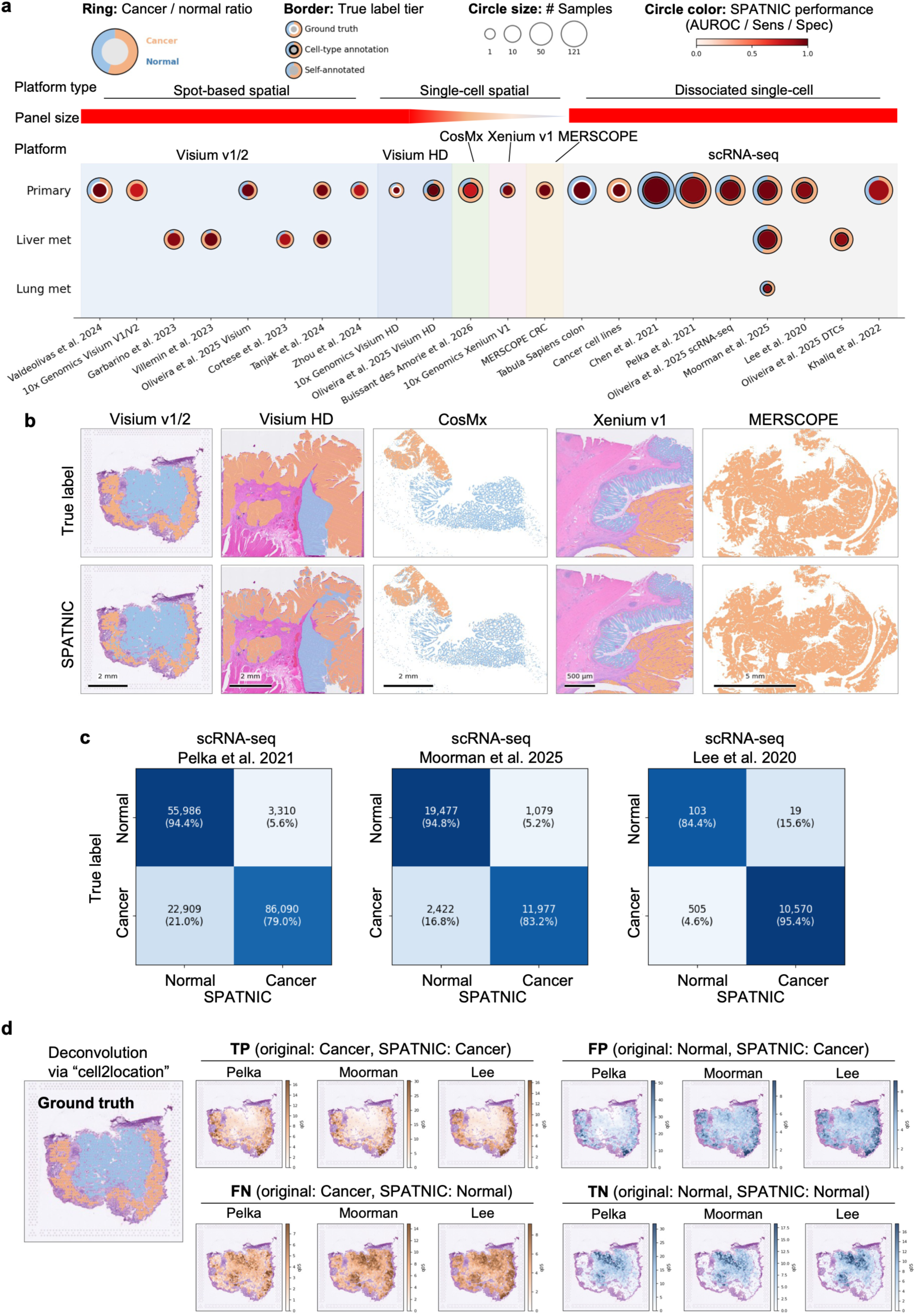
SPATNIC generalizes zero-shot across spatial and single-cell platforms. (a) Applicability- and-performance map of the evaluation datasets, arranged by platform (Visium, Visium HD, CosMx, Xenium V1, MERSCOPE, and scRNA-seq; grouped into spot-based spatial, single-cell spatial, and dissociated single-cell) and by anatomy (primary tumor, liver metastasis, and lung metastasis). Each glyph encodes SPATNIC performance and dataset composition: inner disk color, SPATNIC performance (balanced accuracy where both cancer and normal-epithelial reference labels are available, or sensitivity/specificity where only one class is labeled, with AUROC reported alongside); outer ring, cancer/normal-epithelial composition; glyph size, number of samples; border, reference-label tier (white border, pathologist ground truth; gray border, cell-type annotation; no border, self-annotated). (b) Representative sections across Visium v1/2, Visium HD, CosMx, Xenium V1, and MERSCOPE, each showing the reference label (True label) and the SPATNIC prediction; cancer in orange, normal in blue. (c) SPATNIC predictions compared with the original author annotations on three CRC scRNA-seq atlases (Pelka et al. 2021, Moorman et al. 2025, and Lee et al. 2020), with cells colored by the original annotation (true label) and by the SPATNIC prediction. (d) Spatial deconvolution (cell2location) of the four concordance categories—true positive (TP), true negative (TN), false positive (FP), and false negative (FN)—onto pathologically annotated Visium tissue for each atlas; the matched ground-truth region is shown at left.

SPATNIC maintained its classification ability across all major platform types represented in the collection, although its performance varied across platforms: imaging-based and high-resolution spatial platforms (Xenium V1, MERSCOPE, CosMx, and Visium HD), spot-based spatial platform Visium v1/2, and dissociated single-cell profiling without spatial information (scRNA-seq) (Fig. 3a). The cross-platform performance was not limited to whole-transcriptome assays. SPATNIC achieved AUROCs of 0.99 on Visium v1/2, 0.97–1.00 on Visium HD, and 0.93–0.98 on scRNA-seq, and also transferred to smaller targeted spatial transcriptomic panels, including the ∼6,000-plex CosMx (AUROC 0.75), the 422-gene Xenium V1 (AUROC 0.98), and the 500-gene MERSCOPE (sensitivity 0.99; insufficient annotated normal-epithelial cells for a meaningful AUROC calculation) (Fig. 3a, Fig. S5a, b). In tissues, SPATNIC predictions showed spatial concordance with the available reference annotations for each platform (Fig. 3b); detailed comparisons for Visium and Visium HD are shown in Fig. S6a, b. To emulate smaller panels, we restricted Xenium V1 expression to subsets corresponding to genes shared with the Visium HD (407 genes), Visium v1/2 (419 genes), and Xenium 5K (181 genes overlapping the 3,000 training genes) panels. The classification performance remained high with approximately 400 genes but declined when only the 181-gene Xenium 5K overlap was retained (Fig. S6c). As an orthogonal evaluation using single-class reference datasets, SPATNIC achieved high sensitivity in CRC cell lines^36^ and high specificity for normal colonic tissue (Tabula Sapiens)^37^ (Fig. S6d). Taken together, these results demonstrate cross-platform generalization without platform-specific retraining and robustness to substantial reductions in available gene features.

### Discordant SPATNIC calls in CRC scRNA-seq mark distinct cell populations

Next, we applied SPATNIC to large-scale CRC tissue scRNA-seq, which captured diverse cellular populations within tumors and their microenvironments. Using three independent atlases^19,20,21^, we compared the SPATNIC labels with the original author annotations (Fig. 3c). Cross-tabulation of the two yielded the four expected categories defined relative to the original annotations for convenience: true positives (TP; annotated cancer, predicted cancer), true negatives (TN; annotated normal, predicted normal), FPs (annotated normal, predicted cancer), and FNs (annotated cancer, predicted normal). Most cells were concordant—79.0–95.4% of annotated cancer cells and 84.4–94.8% of annotated normal cells received matching SPATNIC labels across datasets; however, a substantial minority of cells fell into the discordant FP and FN categories (Fig. 3c).

To obtain independent spatial support for these assignments, we examined the localization of the four categories in the tissue. We deconvoluted all four categories into independent, pathologically annotated Visium specimens using cell2location^38^ and quantified their spatial enrichment (Fig. 3d and Fig. S7a). As expected, TP and TN cells were enriched in malignant and normal tissue regions, respectively. Despite their original annotations, FP cells were enriched in tumor regions, whereas FN cells were enriched in normal regions. Moreover, pseudo-bulk signature scoring showed that FP cells more closely resembled the cancer (TP) group, whereas FN cells more closely resembled the normal (TN) group across stemness, differentiation, and proliferation axes (Fig. S7b). Taken together, these results suggest that discordant calls may represent biologically distinct cell states that are not fully captured by the original annotations rather than classification errors. To characterize the discordant FP and FN populations, we clustered the combined TP+FP and TN+FN cells. Rather than scattering across the embedding, the discordant cells formed discrete clusters: an FP-specific cluster (cluster 12; n = 398) and an FN-specific cluster (clusters 1, 4, and 17; n = 1,652) (Fig. S8a and b), supporting the interpretation that these populations represent coherent cell states rather than random classification errors.

In this study, we focused on FP-specific clusters. Marker analysis revealed a distinct transcriptional signature, these cells co-expressed *POU2F3*, the master transcriptional regulator of the tuft-cell lineage^39^, together with the intestinal stem cell and cancer-associated marker *ASCL2* and canonical tuft markers *TRPM5* and *SH2D6*^40^ (Fig. 4a). We provisionally refer to this population hereafter as tuft-like cancer stem cells. Although these cells shared a tuft lineage transcriptional program with normal tuft cells, they were transcriptionally distinct and separate from normal tuft cells in the expression space (Fig. S9a).

**Figure 4.**
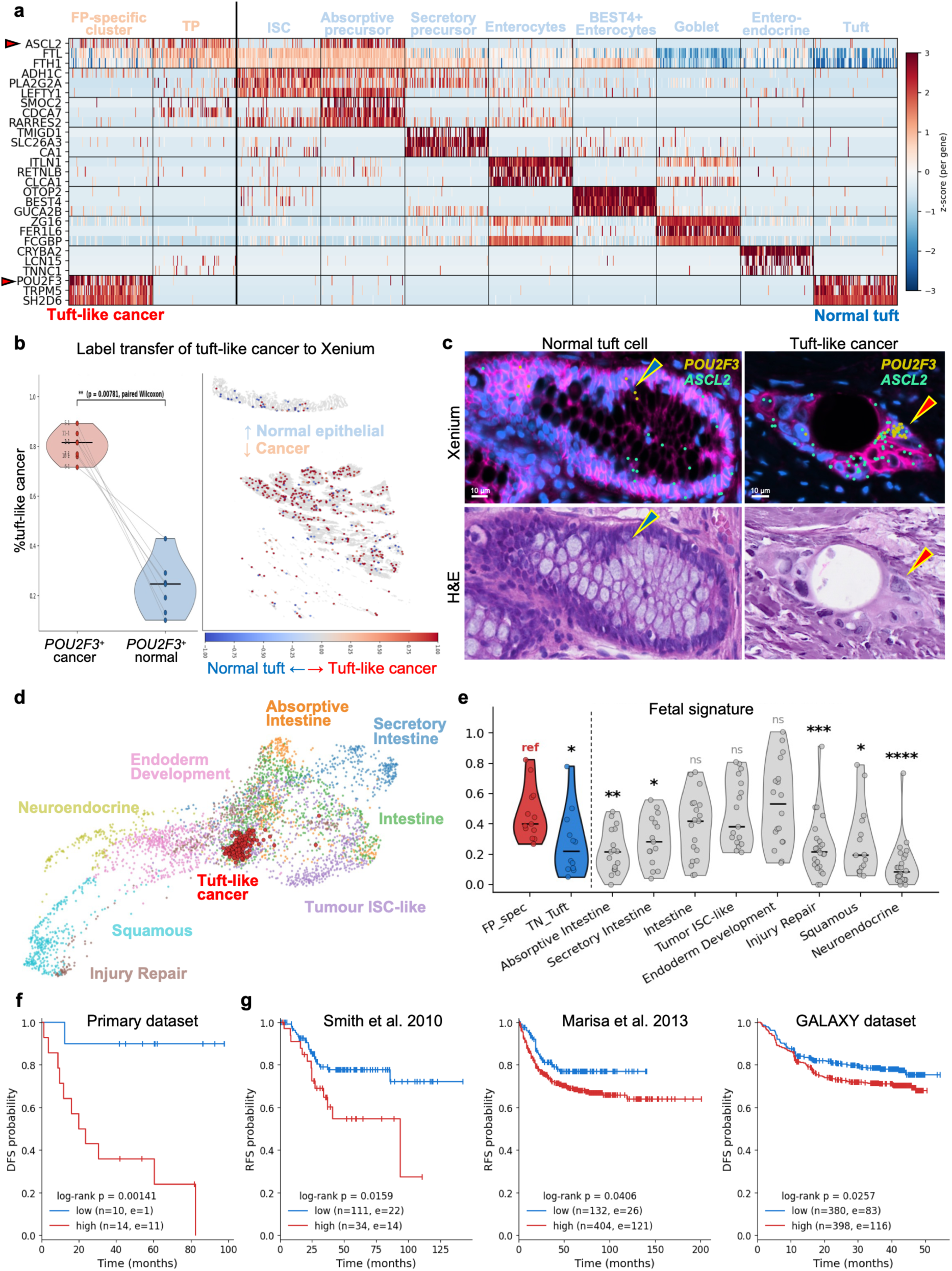
SPATNIC recovers a *POU2F3*⁺ tuft-like cancer stem cell population associated with relapse. (a) Single-cell expression heatmap of marker genes across cell subsets, including the false-positive-specific (FP-specific) cluster, true-positive (TP) cancer cells, and normal epithelial lineages (ISC, absorptive/secretory precursors, enterocytes, *BEST4*⁺ enterocytes, goblet, enteroendocrine, tuft). *POU2F3*, *TRPM5*, and *SH2D6* mark a tuft-like cancer stem cell population distinct from normal tuft cells; color encodes z-scored expression. (b) Quantification of tuft-like cancer stem cell fraction in *POU2F3*⁺ cancer versus *POU2F3*⁺ normal cells (left; paired test) and spatial label transfer of the tuft-like cancer stem cell signature onto Xenium data (right). (c) Validation by Xenium and matched H&E showing *POU2F3* and *ASCL2* co-expression in normal epithelial versus cancer tissue; arrowheads mark tuft-like cells. Scale bar, 10 μm. (d) UMAP of cancer cell states; the tuft-like cancer stem cell population is highlighted among ISC-like, secretory/squamous/neuroendocrine, and injury-repair/endoderm-development programs. (e) Fetal-signature scores across cell subsets relative to the FP-specific cluster reference (ref); significance indicated (ns, not significant; *P < 0.05, **P < 0.01, ***P < 0.001, ****P < 0.0001). (f, g) Kaplan–Meier disease-free/relapse-free survival in the primary dataset, stratified by tuft-like cancer stem cell abundance (f), and in external cohorts (Smith et al. 2010, Marisa et al. 2013, GALAXY), stratified by the tuft-like cancer stem cell signature (high vs low) (g); log-rank P values and group sizes (n, events e) are shown.

Next, we sought independent in situ evidence for this population by using our external Xenium cohort. Projection of the tuft-like cancer stem cell signature onto this independent cohort showed preferential localization to tumors rather than normal epithelial regions (P = 0.0078, paired Wilcoxon; Fig. 4b). At single-molecule resolution, Xenium revealed cells co-expressing *POU2F3* and *ASCL2* within pathologist-annotated tumor regions, a combination observed in malignant regions but not detected in normal tuft cells in this cohort; matched H&E staining showed these cells within tumor-budding regions, where they exhibited a distinctive rounded morphology (Fig. 4c). These observations, obtained directly from tissues on an independent platform, support the interpretation that *POU2F3*⁺ *ASCL2*⁺ tuft-like cancer stem cells belong to the malignant compartment.

We next examined where tuft-like cancer stem cells sit within the landscape of malignant cell states, using the cancer cell-state annotations from Moorman et al.^20^ On the resulting UMAP, tuft-like cancer stem cells occupied an intermediate position between canonical differentiation states (secretory intestine, intestine) and non-canonical states (squamous, neuroendocrine), consistent with a transcriptional state intermediate between distinct differentiation programs (Fig. 4d). This positioning also resembled a fetal progenitor–like state, which Moorman et al.^20^ proposed as a convergent hub through which CRC cells access non-canonical differentiation trajectories. Supporting this, tuft-like cancer stem cells showed significantly higher fetal intestinal signature scores than both the canonical and the non-canonical states (Fig. 4e), consistent with a fetal-like developmental reprogramming phenotype. To define the molecular program underlying this population, we performed a differential expression analysis between the FP-specific cluster and normal tuft cells, which showed increased expression of *ASCL2*, *IGFBP2*, and *HMGB3* in tuft-like cancer stem cells (Fig. S9b). The selected discriminative genes, including *TEAD2*, *GDF11*, and *PTK7*, were specifically enriched in tuft-like cancer stem cells (Fig. S9c), and pathway mapping highlighted enrichment of Hippo-, Wnt-, and TGF-β-related transcriptional programs, including the YAP/TEAD axis, relative to normal tuft cells (Fig. S9d).

Finally, we tested whether tuft-like cancer stem cells were associated with clinical outcomes. In our primary cohort, the abundance of tuft-like cancer stem cells stratified patients by outcome, with a high tuft-like cancer stem cell abundance associated with significantly worse disease-free survival (log-rank P = 0.00141; Fig. 4f). The same stratification using a transcriptional tuft-like cancer stem cell signature rather than cell abundance showed an association in this cohort (log-rank P = 0.00334; Fig. S10a), indicating that the signature-based readout recapitulated the survival association. We then tested whether this signature was associated with outcomes in external bulk-profiled cohorts. The tuft-like cancer stem cell signature stratified disease-free or relapse-free survival across three independent cohorts: Smith et al.^41^ (P = 0.0159), Marisa et al.^42^ (P = 0.0406), and the GALAXY dataset^43^ (P = 0.0257) (Fig. 4g) and showed a consistent trend in the TCGA-COAD/READ cohort^44^ (P = 0.0979, Fig. S10b).

### SPATNIC enables patient-specific screening of ADC target genes in CRC

ADCs deliver a cytotoxic payload to cells expressing a surface target. Their therapeutic index is influenced, in part, by the degree to which target expression is enriched in cancer relative to normal tissue^45^. Because SPATNIC separates cancer from normal epithelium directly in tissues, it can quantify cancer-versus-normal differences in transcript abundance for each patient. We applied SPATNIC to a cohort of patients with locally advanced rectal cancer (LARC) profiled using Visium HD at pre-treatment biopsy and resection after chemoradiotherapy (CRT) (Fig. 5a). For each patient and time point, SPATNIC partitioned the epithelium into cancer (Epi-Tumor) and normal (Epi-Normal) compartments, and we computed the differential expression of a curated panel of ADC target candidates, spanning approved agents, clinical-stage candidates, and undeveloped surface antigens^45^ between the two compartments, stratified by treatment response (non-response, NR; major pathologic response, MPR; pathologic complete response, pCR) (Fig. 5b; the full 87-antigen panel is shown in Fig. S11). Several targets showed cancer-enriched expression that varied across patients and time points. For example, ADC-related antigens, such as *ERBB2/HER2* and *CEACAM5* showed tumor-enriched expression in several pretreatment samples but did not show clear tumor expression in post-CRT specimens. In contrast, *CEACAM6* and *DPEP1* showed tumor-enriched expression in four and two of the seven post-CRT NR cases, respectively (Fig. 5c, d). This analysis revealed that the cancer-specific expression of ADC target candidates varied substantially across patients as well as before and after CRT (Fig. 5b). These findings suggest that both inter-patient heterogeneity and temporal changes in target expression may be relevant when prioritizing ADC candidates for further evaluation. In addition, in several patients, most tumor-specific candidates were undeveloped antigens rather than approved targets. These findings indicate that this screening may support the prioritization of established ADC targets and the nomination of new candidate targets for further validation.

**Figure 5.**
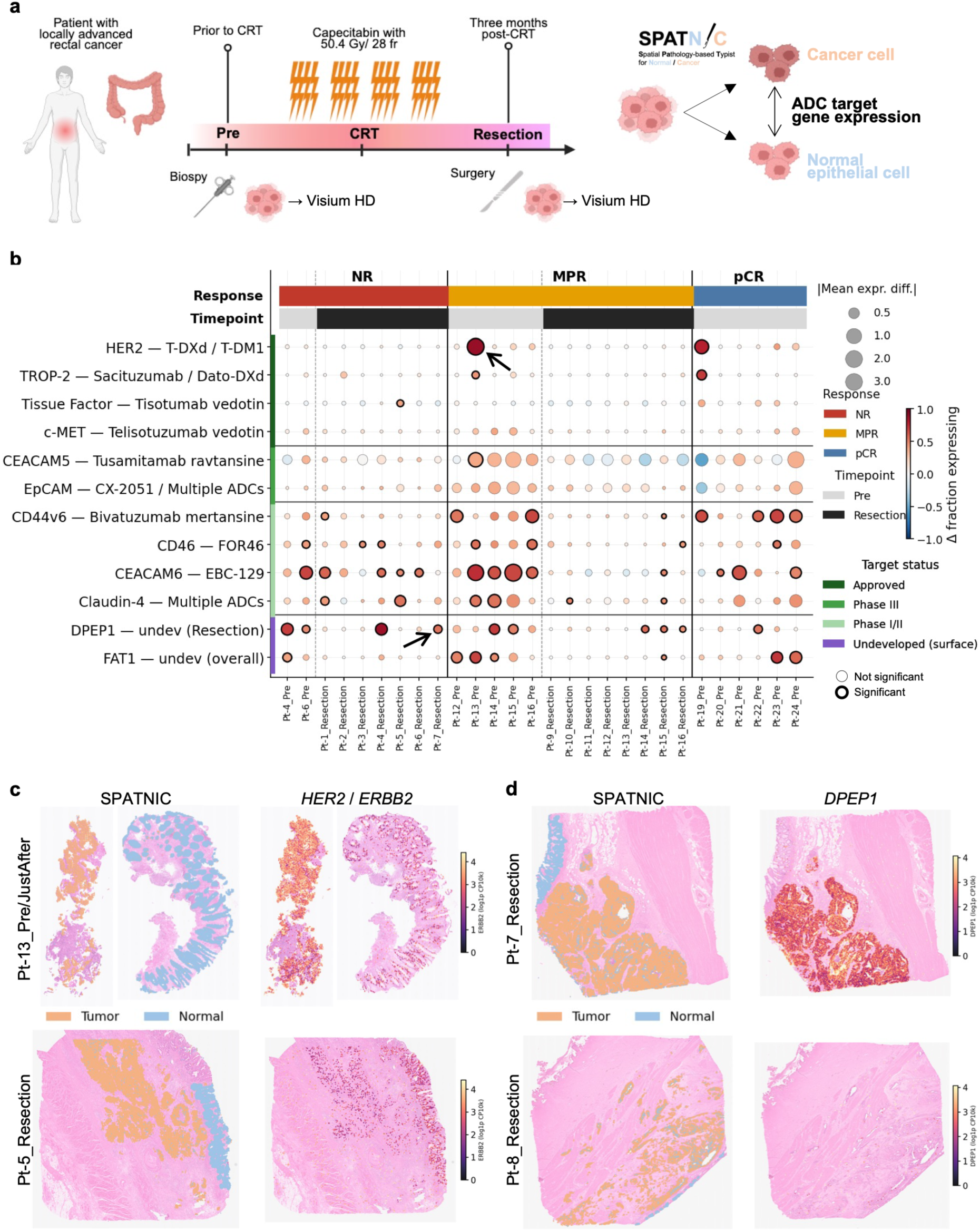
SPATNIC enables patient-specific screening of ADC target genes in colorectal cancer. (a) Study design: locally advanced rectal cancer (LARC) patients profiled by Visium HD at biopsy (pre-CRT) and resection (post-CRT, after capecitabine with 50.4 Gy/28 fr). SPATNIC-classified cancer versus normal epithelial cells are compared for ADC target gene expression. (b) Dot plot of differential ADC-target expression (cancer vs normal epithelium) across patients, grouped by response (NR, MPR, pCR) and timepoint (pre, resection). Dot color encodes the difference in the fraction of expressing bins and dot size the mean expression difference (cancer vs normal); bold outlines denote significance (FDR < 0.05 and AUROC ≥ 0.7); target status (approved / phase III / phase I-II / undeveloped surface) is annotated at left with corresponding agents. (c, d) Spatial validation of representative targets: SPATNIC tumor/normal classification alongside expression of HER2/*ERBB2* (c) and *DPEP1* (d) in selected patients; tumor in orange, normal in blue, with per-gene expression color scales.

To assess whether these differential expression calls reflected genuine spatial localization rather than compartment-averaging artifacts, we examined representative targets in situ. For the approved target HER2/*ERBB2*, the SPATNIC-defined tumor and normal regions overlaid with the target expression showed enrichment of *ERBB2* transcripts in the SPATNIC-classified cancer regions in representative patients (Fig. 5c). For the undeveloped antigen *DPEP1*, the resection sample of Pt-7, a tumor for which existing ADC targets offered no clear candidates, showed *DPEP1* enrichment within SPATNIC-classified cancer regions, illustrating how the screen can nominate additional candidate targets when established ADC targets show limited tumor enrichment (Fig. 5d). These results illustrate how patient-specific ADC target candidates can be identified from spatial data using SPATNIC to separate cancer cell from normal epithelium, providing a hypothesis-generating framework for prioritizing targets that warrant further protein level and functional validation.

## Discussion

We developed SPATNIC, a variational autoencoder that classifies individual cells as cancer or normal based on their gene expression. SPATNIC accurately identifies cancer cells across primary colorectal tumors and their liver and lung metastases, matches or exceeds the performance of benchmarked classifiers while requiring fewer computational resources, and generalizes across single-cell and spatial platforms without platform-specific retraining. Beyond classification, it identified a rare cancer cell subset from public data and nominated patient-specific candidate ADC targets, illustrating how accurate cancer/normal classification can provide a foundation for downstream biological and translational analyses.

Several design choices distinguish SPATNIC from preexisting classifiers. Current tools rely either on inferred copy-number variations or on relatively expression-based classification models; scMalignantFinder^9^ and ikarus^10^ are built on logistic regression over curated signatures or differentially expressed genes, and CancerFinder^8^ applies domain generalization to a feed-forward network. However, the benchmarked methods were developed using labels derived primarily from prior annotations or expression-based criteria, rather than matched histopathology. SPATNIC differs firstly in its training labels; each training cell is labeled as cancer or normal by a pathologist reviewing the matched H&E images. As labels are grounded in histology rather than expression, they reduce potential circularity that arises when labels and features are derived from the same data. This design may contribute to the robust separation achieved by SPATNIC. From a modeling perspective, the heavy tails of the Student-*t* likelihood reduce the influence of extreme and highly variable counts in sparse single-cell and spatial data, which may help the learned representation rely less on platform-specific variation.

The portability of SPATNIC across platforms addresses a practical barrier in a field where laboratories use diverse transcriptomic technologies. Some existing tools, including scMalignantFinder^9^, also report transfer from scRNA-seq training to spatial data, such as Visium, what distinguishes SPATNIC is that the corresponding organ-specific model generalized without platform-specific retraining across a broader span of modalities—Visium, Visium HD, CosMx, Xenium, MERSCOPE, and scRNA-seq—including targeted imaging panels on which copy-number- and signature-based approaches are not designed to operate. As new platforms such as Atera^46^ continue to emerge, models that can accommodate new assays without platform-specific retraining will become increasingly valuable. The cross-platform generalization observed here suggests that SPATNIC can facilitate cancer cell identification as technologies evolve and enable its retrospective application to the growing body of publicly available single-cell and spatial transcriptomic datasets.

The two applications presented illustrate the broad utility of this tool. Applied to publicly available scRNA-seq datasets, SPATNIC identified a rare cancer cell subset co-expressing *POU2F3* and *ASCL2* that had not been identified as malignant in the original annotations. Tuft-like malignant populations have been described as distinct subsets across multiple organs^47^, including *IL17RB*⁺ tuft-like cancer stem cells in the colorectum^48^. Consistent with this, human intestinal tuft cells have recently been shown to express intestinal stem-cell–associated genes, including *ASCL2*, and to act as a damage-induced reserve stem cell pool^49^, providing a normal tissue counterpart to the hybrid tuft stem phenotype observed here. Such noncanonical transcriptional phenotypes may be difficult to capture using conventional annotation or fixed-signature approaches. SPATNIC classified the *POU2F3*⁺*ASCL2*⁺ population as cancer, a prediction supported by independent spatial and histological evidence, suggesting that the model can identify malignant populations that may be missed by conventional annotation. The association of this population within a fetal-like transcriptional state and its transcriptional signature with poor survival across independent cohorts further illustrates how SPATNIC can detect biologically and clinically relevant signals in previously analyzed datasets. Applied to the spatial profiles of treated tumors, SPATNIC partitioned cancer from the normal epithelium and enabled the nomination of patient-specific ADC target candidates, extending cancer cell classification to translational hypothesis generation. Both applications used pretrained SPATNIC models without task-specific retraining.

Our study has several limitations. First, SPATNIC is currently developed and validated only for CRC, including primary tumors, and liver and lung metastases. However, whether the same approach can be extended to other cancer types remains unclear. Secondly, the downstream biological and translational findings reported herein are both exploratory and associative. ADC target candidates are nominated on the basis of differential expression in tissue and require protein-level, experimental, and clinical validation before their therapeutic relevance can be established; however, the causal contribution of the tuft-like cancer stem cell population to disease recurrence remains unproven.

Despite these limitations, SPATNIC provides an accurate, interpretable, and cross-platform foundation for the identification of cancer cells. As single-cell and spatial datasets continue to be accumulated across an expanding range of technologies, a classifier trained on histology-grounded labels capable of operating across modalities offers a scalable means of revisiting existing data and generating reliable cancer-versus-normal cell assignments that support a wide range of downstream analyses.

## Methods

### Data collection and ethics

CRC specimens were profiled in-house using 10x Genomics Xenium *in situ* gene expression. The primary training resource comprised 52 rectal cancer samples. Independent CRC cohorts, external primary cancer, liver metastasis, and lung metastasis, were profiled using the same Xenium workflow and were used for evaluation only (Table 1). Human tissue collection and use were approved by the institutional review board (IRB) of the National Cancer Center, and written informed consent was obtained from all participants. Data from the LARC cohort reanalyzed as shown in Fig. 5 were collected at the National Cancer Center Hospital East under IRB protocol 2018-101: tumor and adjacent normal tissues were obtained from 24 patients with LARC who underwent surgical resection, sampled at two standard timepoints—pretreatment (Pre) and at surgical resection (Resection)—with additional within-2-weeks post-chemoradiotherapy (JustAfter) biopsies from four patients; Pre and JustAfter samples were collected by biopsy forceps and Resection samples by surgical excision.

Public datasets reanalyzed in this study were used under their original licenses: the colorectal scRNA-seq atlases of Pelka et al.^19^, Moorman et al.^20^, Lee et al.^21^, Chen et al.^34^ (colorectal polyp atlas), and Khaliq et al.^35^; the spatial cohorts of Valdeolivas et al. (Visium) and Oliveira et al. (Visium HD; GEO GSE280318); the 10x Genomics public Xenium and Vizgen MERSCOPE colorectal panels and the NanoString CosMx colorectal cancer atlas of Buissant des Amorie et al.^14^; the primary and liver-metastasis Visium cohorts of Zhou et al.^32^, Tanjak et al.^33^, Villemin et al.^29^, Garbarino et al.^30^, and Cortese et al.^31^; the bulk cohorts GSE39582 (Marisa et al.^42^), GSE17536 (Smith et al.^41^), the GALAXY/CIRCULATE-Japan cohort^43^, and TCGA-COAD/READ^44^. The LARC Visium HD cohort analyzed as shown in Fig. 5 is the chemoradiotherapy cohort of Sakai et al.^22^

### Tissue processing and spatial transcriptomics

In situ spatial transcriptomic profiling was performed on a Xenium platform (10x Genomics) according to the manufacturer’s instructions and was optimized for formalin-fixed paraffin-embedded (FFPE) tissue. FFPE sections (5 μm thick) were mounted on Xenium-compatible slides and, after deparaffinization, subjected to antigen retrieval and protease treatment. Gene expression was profiled using the Human Gene Expression Panel (5,001 genes). To ensure comparability across samples profiled with the default panel alone or with an additional add-on panel, analyses were restricted to the 5,001 genes shared between the panels. After signal amplification and imaging, raw data were processed using the Xenium Analyzer pipeline, and single-cell segmentation and transcript quantification were performed to generate spatially resolved gene expression matrices. The single-cell expression matrix, cell centroids in microns, and DAPI morphology images produced by the Xenium Onboard Analysis were used directly. All other spatial and bulk datasets were obtained from public repositories (see the Data collection and ethics section) and were not generated in this study.

Spatial transcriptomic profiling of the LARC cohort was performed on adjacent serial sections using the Visium HD Spatial Gene Expression assay (10x Genomics) according to the manufacturer’s instructions for FFPE samples. FFPE sections were deparaffinized, stained, and imaged to record tissue morphology, and then processed for probe hybridization and library preparation to generate spatially barcoded gene expression libraries, which were sequenced on an Illumina platform. Raw sequencing data were processed using Space Ranger (10x Genomics) to generate spatial gene expression count matrices aligned with the GRCh38 reference genome.

### Pathologist annotation and single-cell-level ground-truth generation from H&E

The cancer/normal ground truth was assigned to individual Xenium cells against matched H&E images. Cells were first clustered with the standard Xenium graph-based clustering, and epithelial clusters were identified from the per-cluster mean log-normalized expression of curated marker panels (an epithelial score): epithelial-general (*EPCAM*, *CDH1*, *KRT20*); colon-tumor (*CDX2*, *MKI67*, *MYC*, *LGR5*, *AXIN2*); colon-normal (*CA2*, *GUCA2B*, *SLC26A3*, *AQP8*, *MUC2*, *MUC4*, *CLCA1*); and, for lung metastases, alveolar (*AGER*, *AQP5*, *HOPX*, *PDPN*, *NKX2-1*, *LAMP3*, *RTKN2*), and bronchial (*TP63*, *FOXJ1*, *MUC5B*, *KRT5*) panels. Clusters scoring above the across-cluster mean were proposed as epithelial, and a per-cluster z-scored tumor-minus-normal marker difference gave an initial tumor/normal call. A pathologist then reviewed the matched H&E images and finalized the labels by lasso selection on the spatial coordinates for primary tumors or for metastases by reviewing per-cluster H&E small multiples on the registered H&E images (see H&E registration) and selecting the malignant clusters. Expression-based epithelial identification and the preliminary tumor/normal call were used only to guide and facilitate this review, and the final cancer/normal label of every cell was assigned by a pathologist based on matched histology. Each cell received a Tumor or Normal label, and cells for which a definitive histological assignment could not be made were left unlabeled and excluded. For liver metastases, labels were taken from an existing histology-based annotation, mapping malignant epithelium to tumors, and hepatocytes, cholangiocytes, and hepatic progenitor-like cells to normal tissues. Cancer cells thus correspond to malignant glands, and normal cells to non-neoplastic compartments: normal mucosa in primaries, hepatocytes and cholangiocytes in the liver, and alveolar and bronchial epithelium in the lung (Fig. 1d).

Train/test splits were defined at the sample level: seven external primary samples were held out for testing only, liver metastasis was split as 12 train/3 test (random, seed 42), and lung metastasis was split as 4 train/1 test. The internal primary cohort was split at the cell level (see Training procedure), and patient-level generalization was assessed using the seven-sample external primary cohort and held-out metastasis cohorts.

### Data preprocessing, quality control and normalization

Unless log-normalized, raw counts were normalized using total-count normalization, followed by log1p. For inference, SPATNIC aligns each dataset to its 3,000 training genes. Genes are matched by symbols (falling back to the gene-symbol annotation column when fewer than half match, as for Ensembl ID datasets), reordered to the training order, and missing genes are zero-filled. The matched-gene fraction was reported and a warning was raised below 50% coverage. The aligned expression was standardized per gene using the training mean and standard deviation stored at the model checkpoint. The 3,000 highly variable genes were selected from the training data (Seurat method).

### SPATNIC model architecture

SPATNIC is a heteroscedastic variational autoencoder with a Student-*t* reconstruction likelihood and classification head. The encoder maps the standardized 3,000-gene vector through two 512-unit ReLU layers (dropout 0.2) to a 32-dimensional latent Gaussian sampled using the reparameterization technique. The decoder mirrors this (32 → 512 → 512) and, being heteroscedastic, outputs a per-gene mean and log-variance. A classifier head (32 → 128 → number of classes, dropout 0.2) maps the latent sample to class logits, which are converted to class probabilities by softmax. The model is specified as follows:

Let v denote the log1p-transformed, total-count-normalized expression of a single cell over G = 3,000 training genes. Each gene g is standardized with the training-set mean m_g and standard deviation s_g stored in the model checkpoint; during training only, each standardized feature is independently masked with probability p = 0.2 (denoising), giving the encoder input u:

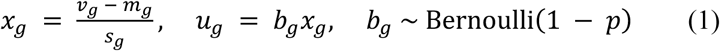

The encoder parameterizes the approximate posterior over the latent variable z of dimension 32, from which a sample is drawn using the reparameterization technique.

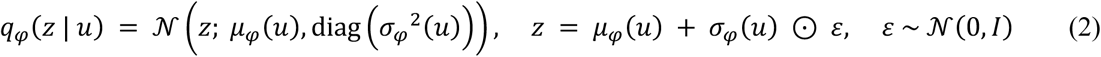

The decoder returns, for each gene, a location and a scale obtained from the clamped log-variance, and reconstructs the unmasked standardized vector x under an independent Student-*t* likelihood with fixed degrees of freedom ν = 5:

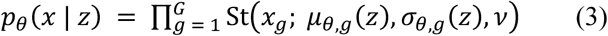

so that the reconstruction loss for one cell is the negative log-likelihood

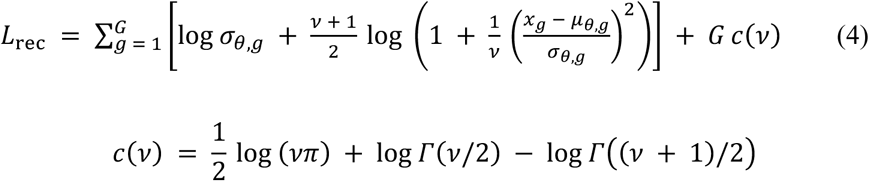

where Γ is the gamma function and c(ν) is constant with respect to the model parameters. As ν → ∞, equation (4) converges to the Gaussian negative log-likelihood; at ν = 5 the squared standardized residual enters logarithmically rather than linearly, so the gradient contributed by any single gene is bounded. This is the sense in which the heavy tails reduce the influence of extreme expression values and highly variable counts pervasive in sparse single-cell and spatial data.

The latent code is regularized toward a standard normal prior by the closed-form Kullback–Leibler divergence, and a classifier head maps the latent sample to class logits ℓ, which are converted to class probabilities by softmax:

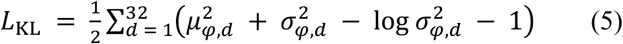

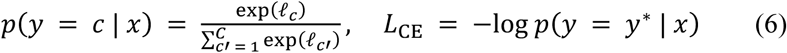

where y* is the pathologist-assigned label of the cell and C is the number of classes. The training objective is the weighted sum.

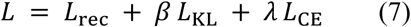

with β = 0.1 throughout, λ = 0 during stage 1 (reconstruction pretraining) and λ = 2.0 during stage 2 (joint training); the optimizer and schedule are given below. During inference, the decoder is skipped, only the encoder and classifier are evaluated, feature masking and dropout are disabled, and the latent code is still drawn stochastically, as in Equation (2), so that the inference path reproduces the classifier input seen during training. The positive-class (tumor) probability from Equation (6) is reported per cell, together with the per-class probabilities.

### Training procedure

The models were trained in two stages. For the primary cancer/normal model, the 52-sample epithelial cells were class-balanced by random subsampling to 244,509 cells per class, split 80/20 into train/test sets (stratified, seed 42), and standardized using statistics computed on the training split. Training used the AdamW optimizer (learning rate 1×10⁻³, weight decay 1×10⁻²), batch size 1,024, gradient-norm clipping at 5.0, and a cosine-annealing learning-rate schedule. Both stages optimize the objective in equation (7): stage 1 (pretraining, 8 epochs) sets λ = 0, and stage 2 (joint, 40 epochs) sets λ = 2.0, with β = 0.1 throughout. The checkpoint with the highest validation accuracy was retained. Class balancing was applied during training and the test sets preserved the native class composition. Each checkpoint bundles the weights, 3,000 gene names, per-gene means and standard deviations, label maps, and model configurations. The second model was trained identically on all cell types to discriminate between cancer cells and all other cells.

### Organ-specific fine-tuning

Liver- and lung-metastasis models were obtained by fine-tuning the primary checkpoint on each organ’s training cells, again in two stages: stage 1 (5 epochs) updates only the classifier head with the encoder and decoder frozen (learning rate 1×10⁻³), and stage 2 (20 epochs) unfreezes all weights and trains jointly (learning rate 5×10⁻⁴, cosine schedule), keeping the same loss weights (β = 0.1, λ = 2.0), optimizer, batch size and an 80/20 stratified cell-level validation split used solely for checkpoint selection. The training cells were class-balanced (up to 100,000 per class for the liver and balanced to the minority class for the lung), and the best-validation checkpoint was retained. To assess transferability, each organ-specific model was applied across all test cohorts. The resulting model × cohort AUC matrix is shown in Tables S2 and S3.

### Strict-confidence (abstention) mode

A strict mode abstains on low-confidence cells. Two thresholds were calibrated on the held-out primary test set so that retained calls meet a target error rate: a lower threshold at the 1st percentile of Tumor scores (≈0.095) and an upper threshold at the 99th percentile of normal scores (≈0.941), so that ≈99% of tumor cells lie at or above the lower threshold and ≈99% of normal cells lie below the upper threshold. Because cells falling between the two thresholds are abstained rather than assigned, these values describe the behavior at each bound and are not equivalent to the conventional binary sensitivity and specificity. The thresholds were calibrated on the held-out primary test set and then applied unchanged to the validation set, for which the strict-mode results are reported (Fig. S2); therefore, the operating points were not selected from the data used to evaluate them. A cell was considered normal below the lower threshold, tumor at or above the upper threshold, and abstained in between these thresholds. Relative to the default 0.5 threshold, this reduced both FN and FP rates, while retaining a subset of cells (Fig. S2a, b), with abstained cells localized to tumor–normal interfaces (Fig. S2c). Unless otherwise stated, all other analyses used the default mode, classifying every cell at a 0.5 threshold.

### Interpretation of discriminative genes

To identify the genes that contributed to this classification (Fig. S1), gene-level attributes were computed for the classifier using GradientSHAP (Captum)^50^. The trained model was wrapped to expose only the class logits, and attributes toward the tumor class were computed for a sample of held-out test cells using a zero baseline in the standardized feature space, which corresponds to the training-set mean of each gene (64 noise samples per cell) and averaged across cells to yield signed per-gene importance—positive values pushing the prediction toward the cancer class and negative values toward the normal class. Leading cancer-class genes (e.g., *SLC2A1*/GLUT1, *ALDOA*, *PGK1*, *VEGFA*, *HMGA1*, *AXIN2*, *SLC40A1*, *FAT1*, *GDF15*) and normal epithelial genes (e.g., *CDX1*, *MUC4*, *MUC5B*, *FCGBP*, *KLF4*, *HOXB13*, *CA12*) are reported. Surface localization of the attributed genes was annotated from Gene Ontology cellular-component terms (via MyGene.info) and UniProt membrane-topology records to flag membrane and extracellular domain genes. Differential expression of representative markers between tumor and normal test cells was confirmed by Wilcoxon rank-sum test with Benjamini–Hochberg correction, and for representative markers (e.g., *SLC2A1*, *MUC4*), at the protein level by immunohistochemistry. The immunohistochemistry images shown in Fig. S1 were obtained from the Human Protein Atlas^51^ (proteinatlas.org).

### Architecture ablation

To attribute robustness to the Student-*t* likelihood, SPATNIC was compared with a Gaussian-likelihood VAE—identical in architecture, latent dimension, schedule and loss weights but replacing the Student-*t* reconstruction with a fixed-variance Gaussian likelihood, for which the loss in equation (4) reduces to ½ ∑_g (x_g − μ_θ,g)² up to an additive constant (mean-squared error)—and with supervised baselines trained on the same standardized 3,000-gene features: a multilayer perceptron, logistic regression (max 1,000 iterations), a random forest (300 trees, seed 42), and XGBoost (400 trees, maximum depth 6, learning rate 0.1, histogram tree method, seed 42). All models were evaluated on the internal primary held-out test set and external primary cohort (domain shift), and the results are shown in Fig. S3, and Tables S6 and S7.

### Benchmarking against existing classifiers

SPATNIC was benchmarked against CancerFinder^8^, scMalignantFinder^9^ and ikarus^10^, each in two configurations: with its published pre-trained weights and after retraining on our CRC data (the same 391,214-cell balanced primary training set over the 3,000 training genes). Each tool was run through its published interface with the required preprocessing: CancerFinder on the standardized matrix via its inference script, scMalignantFinder with its ordered feature set (missing features zero-filled), and ikarus with its signature set filtered to the genes present and processed in chunks. SPATNIC was applied with the released cancer/normal checkpoint without retraining. Apart from the change in the training data, each baseline was run using the feature selection procedure and hyperparameters specified in its published implementation, and no additional hyperparameter search was performed for any method. All methods used a 0.5 threshold for hard labels, and the performance was summarized pooled over all cells and per sample.

### Metrics, statistics, and reproducibility

For each prediction, we computed the AUROC, AUPRC, Matthews correlation coefficient, F1, and—from the confusion matrix at a threshold of 0.5—sensitivity, specificity, and balanced accuracy (positive class = tumor). Pooled-cell metrics are reported with 95% confidence intervals from 1,000 bootstrap resamples (seed 42; Tables 2, S2–S6). Per-sample metrics are reported as mean ± standard deviation across samples, excluding samples with too few cells or a single class present (Tables S1, S4). SPATNIC was compared with each baseline using a two-sided paired Wilcoxon signed-rank test on per-sample AUPRC, with P < 0.05 considered significant. Random seeds were fixed at 42 throughout.

### Compute and memory profiling

Inference time and peak host memory were profiled across cell numbers from 1×10³ to 1×10⁸ for SPATNIC and the baseline tools. Up to ≈10⁶ cells, real test data were resampled; for larger sizes, synthetic sparse matrices matching the gene panel and a realistic sparsity were generated in chunks. The wall-clock inference times (excluding I/O) and peak resident set sizes were recorded. For every method, profiling used the execution mode (CPU or GPU) supported by its published implementation; therefore, the absolute runtimes reflect each tool as a distributed rather than a hardware-matched comparison. The profiling hardware used was an Intel Core i9-14900KF (32 threads) with 62 GB of RAM and an NVIDIA GeForce RTX 4090 (24 GB) operating under Linux (Fig. 2b, c).

### Cross-platform and pseudo-panel evaluation

The primary model was applied without any retraining or fine-tuning to CRC datasets spanning Visium (Valdeolivas et al.), Visium HD (Oliveira et al., GSE280318, and the 10x public Human Colon Cancer dataset), Xenium V1 (10x 422-gene CRC panel), CosMx (NanoString ≈6,000-plex colorectal cancer atlas), MERSCOPE (Vizgen 500-gene panel) and scRNA-seq. Each dataset was normalized as above and classified directly; for spatial cohorts with pathologist annotation, predictions were compared with the histology-based reference annotations (Visium spot annotations; Visium HD 16-µm bin annotations from Zenodo record 11402686, mapping neoplasm to Tumor and non-neoplastic epithelium to Normal). For the CosMx cohort, no matched pathologist annotation was available; the published author cell-type annotations were used as the reference, mapping malignant epithelial labels to Tumor and non-neoplastic epithelial labels to Normal. To assess robustness to gene-panel content, pseudo-panel datasets were built by restricting the test expression to the genes overlapping each target panel and zero-filling the rest, then re-classifying; the overlaps with the 3,000 training genes (Visium HD ≈407, Visium V1/2 ≈419, Xenium 5K ≈181 genes) were compared against random gene subsets of matched size. As expected, the model was applied to CRC cancer cell lines (all expected Tumor) and to normal colons from healthy donors (Tabula Sapiens large intestine; all expected Normal). SPATNIC was compared with the retrained baselines across Visium and scRNA-seq (Fig. S5).

### scRNA-seq application, TP/TN/FP/FN definition and deconvolution

SPATNIC was applied to three CRC tissue scRNA-seq atlases (Pelka^19^, Moorman^20^, Lee^21^), and the predictions were cross-tabulated with the original author cancer/normal annotations to define four categories: TP (author cancer, predicted cancer), TN (author normal, predicted normal), FP (author normal, predicted cancer), and FN (author cancer, predicted normal). For convenience, these designations are used relative to the original author annotations, and do not imply that these annotations constitute a definitive ground truth. To assess whether discordant calls showed independent evidence consistent with their SPATNIC-assigned identities, the four categories were treated as cell-type labels and deconvoluted into independent pathologist-annotated Visium specimens with cell2location^38^. A negative-binomial regression reference signature was estimated per category (with the standard gene filtering, batched by sample, ≈250 epochs), and spatial abundances were inferred with the cell2location model (N_cells_per_location = 8, detection_alpha = 20, up to 30,000 epochs, on GPU). Per-spot category abundances (posterior 5th percentile) were summarized by pathological regions to quantify spatial enrichment (Fig. 3d, Fig. S7a), and pseudo-bulk signature scoring along the stemness, differentiation, and proliferation axes placed the FP and FN cells relative to the TP and TN groups (Fig. S7b). For additional orthogonal validation, discordant calls were assessed against the inferred copy-number burden, curated marker programs, and concordance to normal-adjacent-tissue pseudobulks, each summarized per patient, and compared using the paired Wilcoxon test.

### Clustering, UMAP, and differential expression

TP + FP and TN + FN cells were pooled and embedded separately: 2,000 highly variable genes, principal component analysis (50 components), Harmony^52^ batch integration across datasets, a k-nearest-neighbor graph (k = 30), UMAP, and Leiden^53^ clustering (resolution 0.8). Discordant cells formed discrete, FP-specific, and FN-specific clusters (Fig. S8). Cluster markers were identified using the Wilcoxon rank-sum test (one-versus-rest test). Differential expression between the FP-specific cluster and normal tuft cells was determined using per-sample pseudobulk aggregation and PyDESeq2^54^ (design ∼group; genes filtered by minimum count; significance at adjusted P < 0.05; and |log2 fold change| > 1). Pathway enrichment used Enrichr^55^ (via gseapy^56^) against the MSigDB Hallmark, KEGG and Reactome gene sets, highlighting Hippo, Wnt and TGF-β (YAP/TEAD) activity (Fig. S9d).

### Tuft-like cancer stem cell signature, label transfer, and survival analysis

The FP-specific population was characterized as tuft-like cancer stem cells by co-expression of *POU2F3*, *ASCL2*, *TRPM5* and *SH2D6*. A transcriptional tuft-like cancer stem cell signature was defined from the one-versus-rest differential expression of the FP-specific cluster (log2 fold change > 1, adjusted P < 0.05, high in-group and low out-group detection). The signature was transferred onto the independent external Xenium cohort (7 primary samples) by training scANVI^57^ on the integrated scRNA-seq reference and predicting on *POU2F3*⁺ epithelial Xenium cells; per-sample FP-specific fractions in tumor versus normal epithelial regions were compared by a one-sided paired Wilcoxon signed-rank test (P = 0.0078; Fig. 4b). In situ co-expression of *POU2F3* and *ASCL2* within cancer tissues with matched H&E images provided orthogonal evidence (Fig. 4c). Tuft-like cancer stem cells were positioned within the malignant cell state landscape using the cancer cell state programs of Moorman et al.^20^ (per-cell module-score argmax over each program’s gene set) and scored for a fetal intestinal signature (Fig. 4d, e).

The clinical relevance was assessed using survival analysis (lifelines)^58^. The endpoints were disease-free survival for the primary and GALAXY/CIRCULATE-Japan cohorts, relapse-free survival for GSE17536 (Smith et al.^41^) and GSE39582 (Marisa et al.^42^), and disease-free interval (DFI) for TCGA-COAD/READ^44^ (TCGA-CDR endpoints; primary tumor samples only). In the primary cohort, patients were stratified either by tuft-like cancer stem cell abundance (fraction of FP-classified cells among tumor cells at resection; disease-free survival, log-rank P = 0.00141; Fig. 4f) or by a transcriptional tuft-like cancer stem cell signature that reproduced the association (P = 0.00334; Fig. S10a). The signature was evaluated in external bulk cohorts. For the microarray cohorts (Smith, Marisa) the signature score was the mean log2 expression of the present signature genes; for the bulk RNA-seq GALAXY cohort, a matched per-sample signature score was used, and for TCGA, the score was a control-corrected gene-set score on log-normalized expression. In every cohort, patients were dichotomized at the Youden optimal cutpoint of the score against the survival event, a data-driven cutpoint reported as such, and compared using the log-rank test (Smith P = 0.0159, Marisa P = 0.0406, GALAXY P = 0.0257; TCGA DFI P = 0.0979; Fig. 4g, Fig. S10b).

### ADC target screening on the LARC Visium HD cohort

The LARC cohort of Sakai et al.^22^—24 patients profiled using Visium HD before CRT at biopsy and after CRT at resection, with response groups defined as nonresponse (NR), major pathologic response (MPR), and pathologic complete response (pCR), were analyzed for ADC target candidates. SPATNIC (tumor/normal model) was applied to all 16-µm bins, and epithelial regions were then defined at the cluster level. Bins were clustered per sample with the Space Ranger graph-based clustering, and for each cluster the fraction of bins with P(Tumor) ≥ 0.9 and the fraction with P(Normal) ≥ 0.9 were computed. Clusters for which these two fractions summed to more than 0.5 were proposed as epithelial and were confirmed by review of the matched H&E results together with the per-cluster SPATNIC composition and EPCAM expression, with manual override where the automatic proposal was inconsistent with the histology. Bins outside the selected epithelial clusters were excluded regardless of their SPATNIC scores. Within the retained epithelial clusters, each bin was then assigned at the default 0.5 threshold: a bin was called Epi-Normal if P(Normal) ≥ 0.5 and Epi-Tumor otherwise. Because the two-class probabilities sum to 1, every bin in a selected epithelial cluster receives one of the two labels; therefore, the cluster-level selection above restricts the comparison to the epithelium. Samples with fewer than 10 Epi-Tumor bins or fewer than 10 Epi-Normal bins were dropped from the within-sample comparison, and samples with fewer than 10 Epi-Tumor bins were dropped from the comparison against the pooled global Epi-Normal reference.

We assembled a manually curated, solid-tumor-scoped panel of 87 candidate antigens (Table S8): 72 antigens targeted by clinically developed ADCs or antibody therapeutics, and 15 undeveloped cell-surface candidates nominated de novo (below). Among the 72 developed target antigens, 58 were addressed by a clinical-stage ADC (6 by an approved ADC, 10 by a phase III ADC, and 42 by a phase I/II ADC), and the remaining 14 were targeted by clinically evaluated non-ADC therapies (antibody, bispecific, or cell therapy) for which the ADC modality is currently preclinical or at the investigational-new-drug stage. Each antigen was annotated using the representative agent and the highest clinical phase. Targets of approved ADCs restricted to hematologic malignancies (e.g., CD33, CD30, and CD22) were excluded because they were not expressed in epithelial tumors, and clinical stage assignments were verified against ClinicalTrials.gov, PubMed, and DOI records; 82 of the 87 antigens were detected in the Visium HD gene panel.

To identify novel surface targets, we screened the SURFY in silico surfaceome^59^ (2,799 high-confidence surface proteins). A gene was considered potentially antibody accessible if its SURFY topology indicated no more than six transmembrane domains and the largest predicted extracellular segment of at least 80 amino acids, excluding multipass transporters and endoplasmic-reticulum–resident proteins. Accessible genes were further required to be undeveloped (absent from the curated developed-ADC list) and epithelial-enriched (mean normalized expression ratio of epithelial to non-epithelial bins of at least two), removing stromal, immune, and other microenvironmental contaminants. The resulting 393 candidates were ranked by depth-robust within-sample percentile metrics into three groups: overall (consistently tumor-specific across Pre and Resection), Pre-high (longitudinally higher tumor expression at Pre), and Resection (tumor-high in most Resection samples), and the top five per group (15 in total) were carried forward.

For each sample, per-gene Tumor-versus-Normal differences were computed between the Epi-Tumor and Epi-Normal bins as the difference in the fraction of bins with nonzero raw counts and the difference in mean total-count-normalized, log1p-transformed expression, defining the dot-plot color and size, respectively. Significance per gene × sample was assessed against both the matched within-sample and global pooled Epi-Normal bins using the area under the ROC curve (Tumor vs. Normal) and a two-sided Mann–Whitney U test (each group capped at 2,000 bins by random subsampling), with Benjamini– Hochberg FDR correction across all dots in a plot. A dot was called significant when FDR < 0.05 and AUROC ≥ 0.7 with higher expression in Epi-Tumor than in Epi-Normal bins; the AUROC effect-size gate was required because the large bin counts saturate p values.

Resection samples from patients with a pathologic complete response, which contained no residual tumor, were excluded from the dot plots. To avoid depth confounding, samples were retained only if their mean Epi-Tumor depth was at least 700 reads per bin, leaving 28 samples (13 Pre, 15 Resection; no JustAfter sample met this threshold). For clarity, the main dot plot (Fig. 5b) shows a curated subset of the panel—all approved and phase III antigens passing an expression filter (maximum per-sample fraction of expressing Epi-Tumor bins ≥ 0.30 for approved and ≥ 0.70 for phase III targets), four well-established phase I/II ADC targets (CD44v6/CD44, CD46, CEACAM6 and claudin-4/CLDN4), and two undeveloped surface candidates (*DPEP1* and *FAT1*); the complete 87-antigen panel is shown in Fig. S11. Candidate prioritization also considered cell-surface accessibility, as defined above, and cancer specificity was visualized in situ by overlaying SPATNIC-defined compartments with the target expression (Fig. 5).

### H&E registration and spatial overlays

H&E whole-slide images were registered to the Xenium DAPI morphology image with VALIS^60^. To keep the registration tractable, each whole-slide image was re-encoded at the pyramid level to a plain RGB image (with optional rotation and bounding-box cropping for multi-section slides); only the first channel of the DAPI image was read and downsampled to a single channel, both of which were then registered (non-rigid, cropped to the DAPI reference) at the thumbnail scale on the CPU, and the H&E was warped into Xenium coordinates. Warped H&E was used for both pathologist annotation overlays and side-by-side prediction figures (Fig. 1d, e). The Tumor and Normal tissues are colored in #F4B183 and #9DC3E6, respectively.

### Software

Analyses were performed using Python 3.10 with scanpy 1.10.4^61^, anndata 0.11.4, numpy 1.26.4, scipy 1.15.1, and pandas 2.2.3. The SPATNIC model, training, and inference used PyTorch 2.8 (CUDA 12.6); and gene attributions used Captum 0.8.0^50^. The baselines and metrics used scikit-learn 1.6.1, XGBoost 3.0.5, and statistics used scipy and statsmodels 0.14.6. Deconvolution and label transfer used cell2location 0.1.5^38^ and scvi-tools 1.3.3^57^; differential expression and enrichment used PyDESeq2 0.5.2^54^ and gseapy 1.1.8^56^; batch integration used harmonypy 0.2.0^52^ with rapids-singlecell 0.11.1; survival analysis used lifelines 0.30.0^58^; gene/protein annotation used mygene 3.2.2; dataset retrieval used GEOparse 2.0.4; and H&E registration used VALIS 1.2.0^60^. SPATNIC is distributed in the spatnic Python package (see code availability).

## Data availability

The Xenium and Visium HD data generated from clinical specimens in this study have been deposited in the Japanese Genotype-phenotype Archive (JGA) under accession code JGAS000945 (dataset JGAD001089), registered with the NBDC Human Database under accession hum0461. In accordance with the NBDC guidelines for human data (policy JGAP000001), these data are available under controlled access, and will be released through the NBDC Human Database upon publication; access requests are reviewed by the NBDC Data Access Committee.

In accordance with collaborative research agreements and intellectual property considerations, public data releases may be subject to a temporary embargo (up to 18 months) in the event of patent filing.

The public datasets reanalyzed in Fig. 3a are as follows: Visium: Valdeolivas et al.^28^ (Zenodo 10.5281/zenodo.7760264) and the 10x Genomics public human colorectal cancer Visium products. Visium HD: Oliveira et al.^13^ (GEO GSE280318) and the 10x Genomics public Human Colon Cancer Visium HD dataset. CosMx: Buissant des Amorie et al.^14^ (Zenodo 10.5281/zenodo.17671259). Xenium V1: 10x Genomics public Human Colon Cancer (patient 5) Xenium dataset. MERSCOPE: Vizgen public FFPE Human Colon Cancer dataset. scRNA-seq: Pelka et al.^19^, Moorman et al.^20^, Lee et al.^21^, Chen et al.^34^ and Oliveira et al.^13^. Liver and lung metastases: Moorman et al.^20^, Villemin et al.^29^ (GEO GSE217414), Garbarino et al.^30^ (GEO GSE206552), Cortese et al.^31^ (GEO GSE254364), Zhou et al.^32^ (GEO GSE236697), Tanjak et al.^33^ (GEO GSE267401) and Khaliq et al.^35^ (Zenodo 10.5281/zenodo.6466249).

The bulk and clinical cohorts reanalyzed in this study are available as follows: GSE39582 (Marisa et al.^42^) and GSE17536 (Smith et al.^41^) from the NCBI Gene Expression Omnibus, TCGA-COAD/READ^44^ from the NCI Genomic Data Commons (https://portal.gdc.cancer.gov), and the GALAXY/CIRCULATE-Japan cohort (Nakamura et al.^43^) under the controlled-access data policy of that study.

## Code availability

The SPATNIC model, together with its training and inference codes, is available as the spatnic Python package at https://github.com/shusakai/spatnic. The analysis scripts used in this study are available at https://github.com/shusakai/spatnic_paper.

## Acknowledgments

We thank Dr. Kazuki Nakai, Dr. Hideki Makinoshima, and Dr. Tohru Kiyono for their valuable discussions during in-house meetings. We thank the staff of the National Cancer Center Hospital East for supporting the clinical sample collection and for their helpful advice, and all patients and their families for contributing to this study.

## Author contributions

S.A.S., S.-I.K. and R.Y. conceived and designed the study. S.A.S. and M.O. developed the software and performed computational analyses. Y.Mor., T.K., N.T., M.K., and S.S. performed the histopathological review and cancer/normal annotation of matched H&E images. S.S. provided clinical specimens and associated clinical information. A.S. and Y.S. generated sequencing data. H.E., R.Yg., E.O., T.F. and H.B. provided the data and resources. M.K., K.T., S.-I.K. and R.Y. supervised the study. S.-I.K. and R.Y. acquired funding. S.A.S. wrote the manuscript. All other authors reviewed and edited the manuscript. All authors have read and approved the final version of this manuscript.

