## Supplementary Fig. 1-11 for "SPATNIC enables H&E-grounded cancer-cell identification across spatial and single-cell transcriptomic modalities in colorectal cancer"

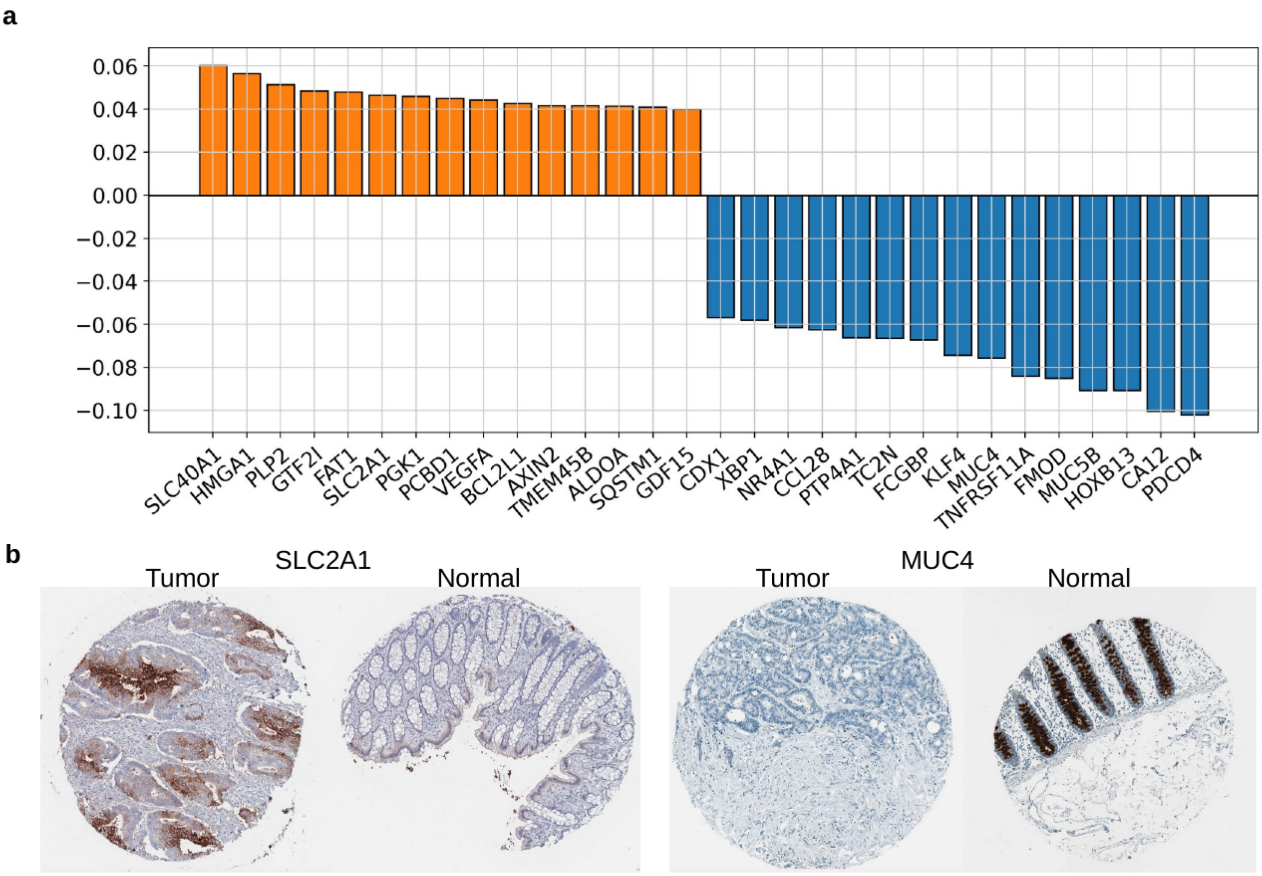


**Supplementary Fig. 1 | Top discriminative genes underlying SPATNIC cancer/normal classification.** (a) Genes ranked by signed contribution to the SPATNIC cancer-versus-normal decision; positive (orange) values favor the cancer class and negative (blue) values favor the normal class. (b) Immunohistochemical validation of two representative discriminative markers, *SLC2A1* (GLUT1) and *MUC4*, in tumor versus normal colorectal tissue, consistent with the model's learned directionality.

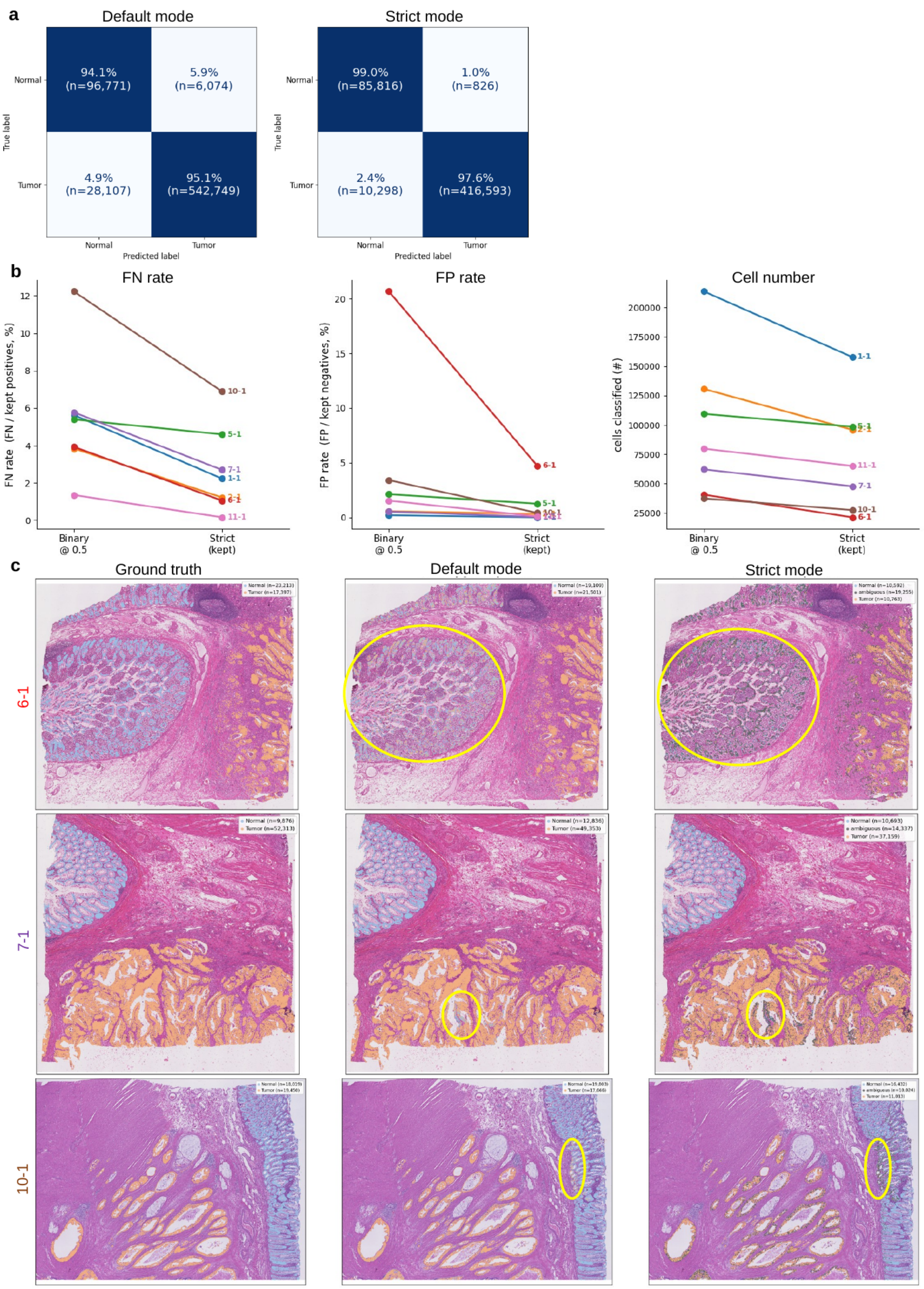


**Supplementary Fig. 2 | A strict-confidence mode reduces false positives by abstaining on ambiguous cells.** (a) Confusion matrices for the primary external test set under default mode (all cells classified) versus strict mode (low-confidence cells withheld); values are row-normalized percentages with cell counts (n). (b) Per-sample false-negative rate, false-positive rate, and number of retained cells comparing binary classification at threshold 0.5 versus strict mode; each line is one section. (c) Representative sections (6-1, 7-1, 10-1) showing H&E ground truth, default-mode, and strict-mode predictions; yellow outlines mark regions where strict mode reassigns histologically ambiguous cells to an "ambiguous" category rather than forcing a tumor/normal call.

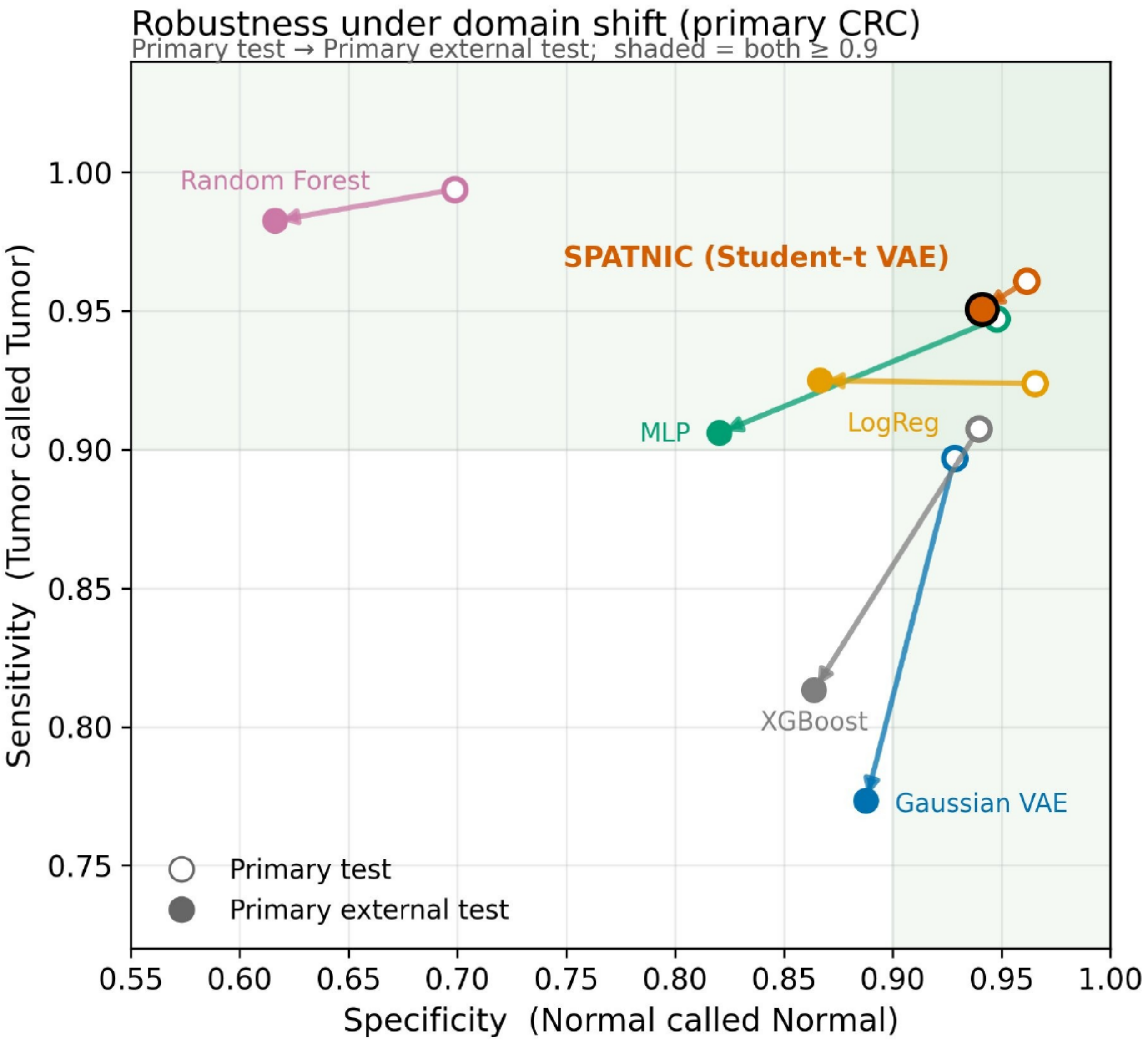


**Supplementary Fig. 3 | Architecture ablation: the Student-*t* reconstruction likelihood underlies SPATNIC robustness under domain shift.** (a) Robustness of SPATNIC and ablated/baseline classifiers under domain shift, shown as sensitivity (tumor recalled as tumor) versus specificity (normal recalled as normal) on the primary test set (open circles) and the primary external test set (filled circles); arrows connect the two for each method, and the shaded region marks both metrics ≥ 0.9. SPATNIC (Student-*t* VAE) is compared with a Gaussian-likelihood VAE that is identical except for the reconstruction likelihood, and with supervised baselines (MLP, logistic regression, random forest, XGBoost) trained on raw features. SPATNIC remains in the high-performance region under domain shift, whereas the Gaussian VAE collapses in sensitivity and the raw-feature baselines lose specificity. Full metrics for internal and external test sets are provided in Supplementary Table S2.

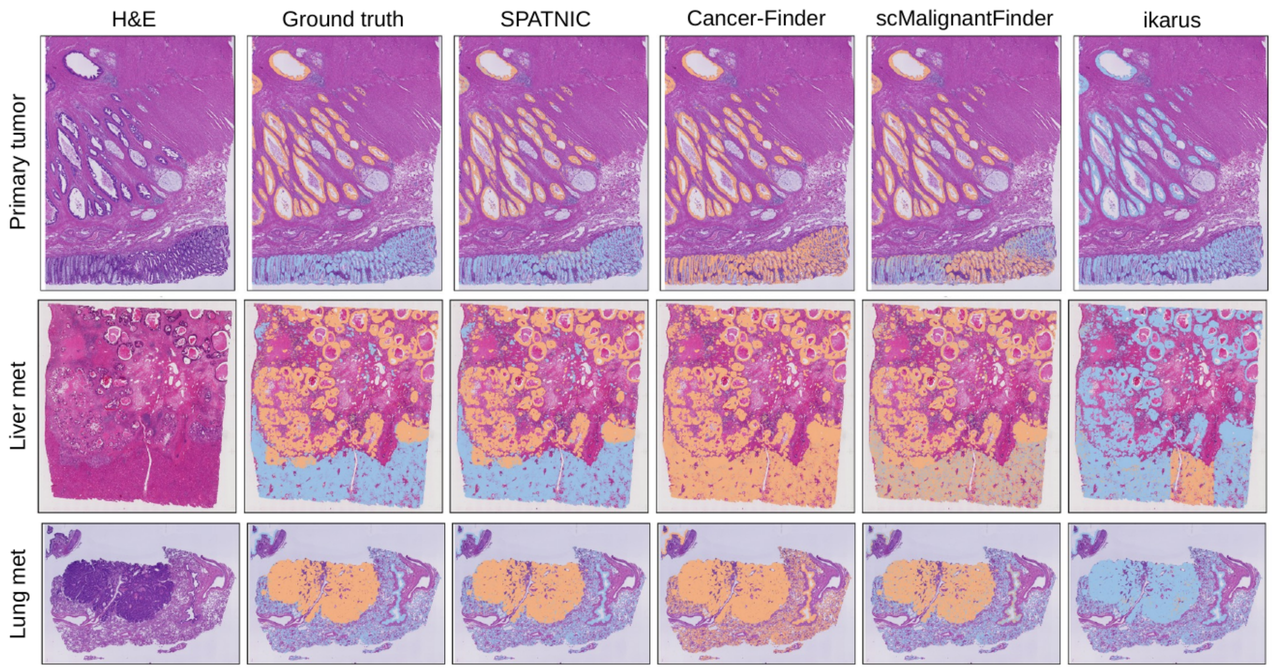


**Supplementary Fig. 4 | Representative spatial classifications for primary tumor and metastases.** Representative sections of a primary tumor, a liver metastasis, and a lung metastasis, showing H&E, pathologist ground-truth annotation, and predictions from SPATNIC, Cancer-Finder, scMalignantFinder, and ikarus (default configurations); cancer in orange, normal in blue.

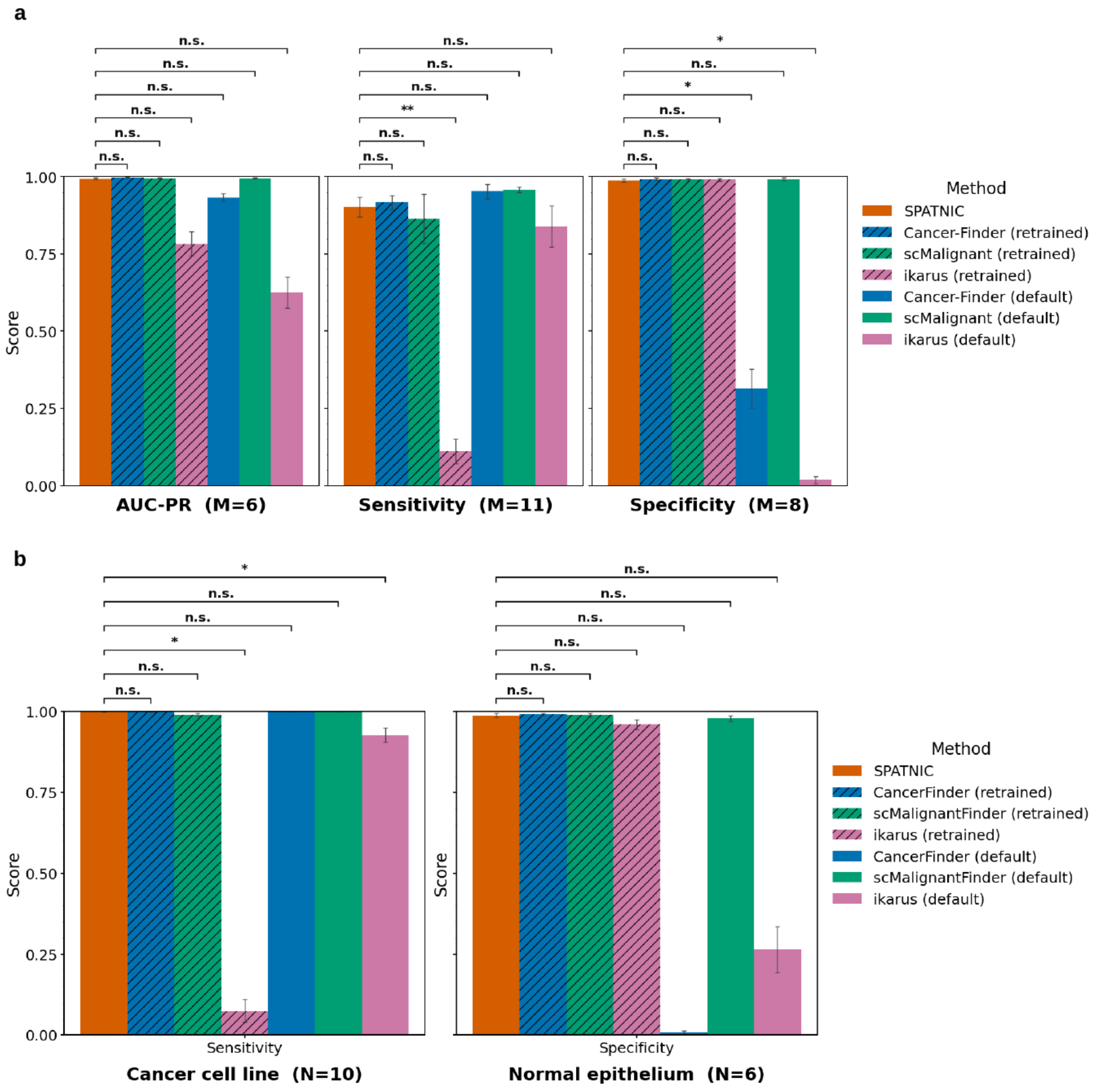


**Supplementary Fig. 5 | SPATNIC matches or exceeds existing tools across Visium and scRNA-seq benchmarks.** (a) AUC-PR (M = 6 datasets), sensitivity (M = 11), and specificity (M = 8) for SPATNIC versus CancerFinder, scMalignantFinder, and ikarus (retrained and default configurations) on Visium data. Bars show mean ± s.d.; significance versus SPATNIC is indicated (n.s., *, **). (b) Sensitivity on cancer cell lines (N = 10) and specificity on normal epithelium (N = 6) for the same methods on scRNA-seq data.

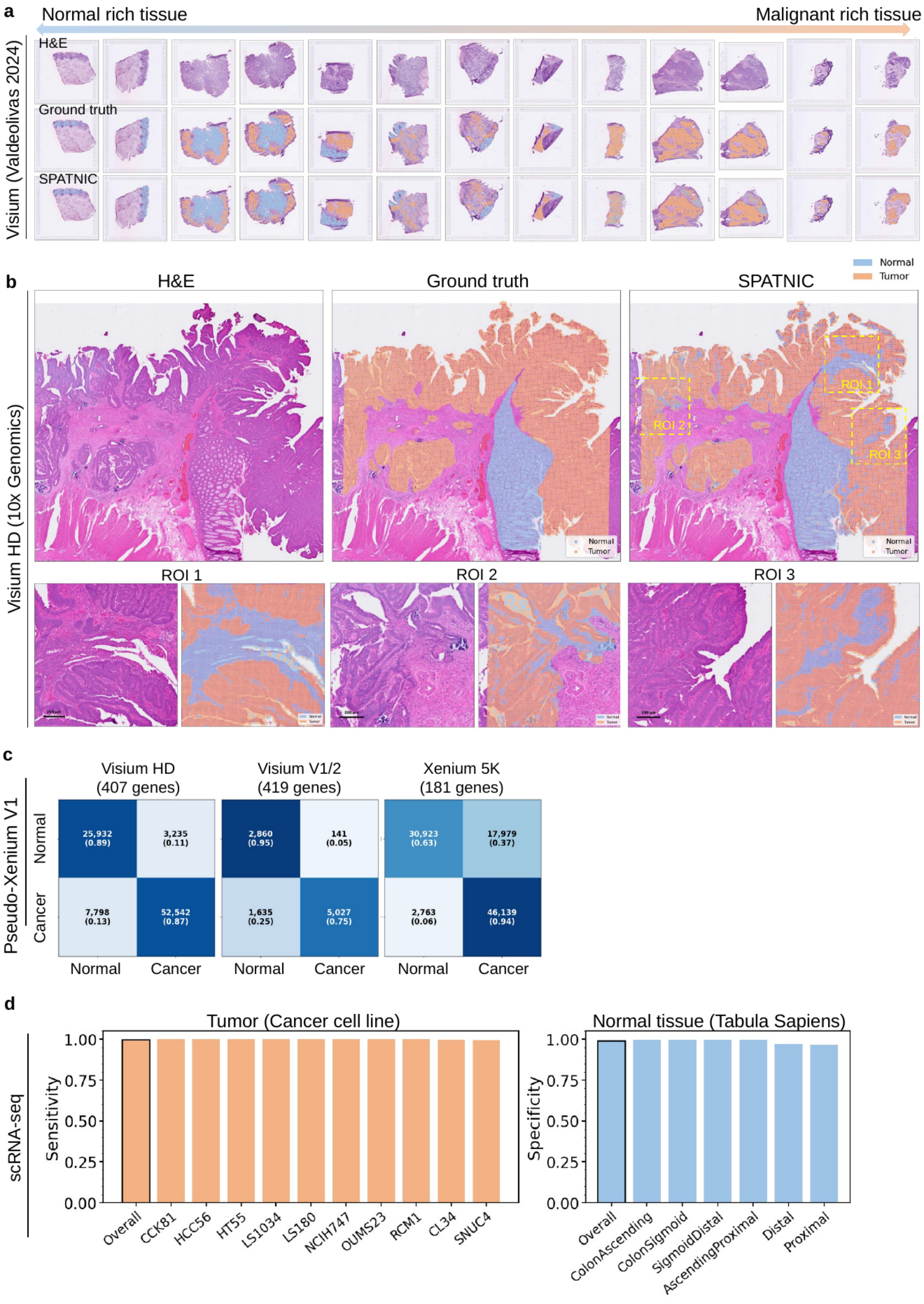


**Supplementary Fig. 6 | SPATNIC generalizes zero-shot to Visium and scRNA-seq.** (c) Pseudo–Xenium V1 evaluation in which expression is restricted to genes overlapping each panel — Visium HD (407 genes), Visium V1/2 (419 genes), and Xenium 5K (181 genes); confusion matrices give counts and row-normalized fractions, illustrating robustness as panel overlap decreases and the expected drop with the smallest panel.

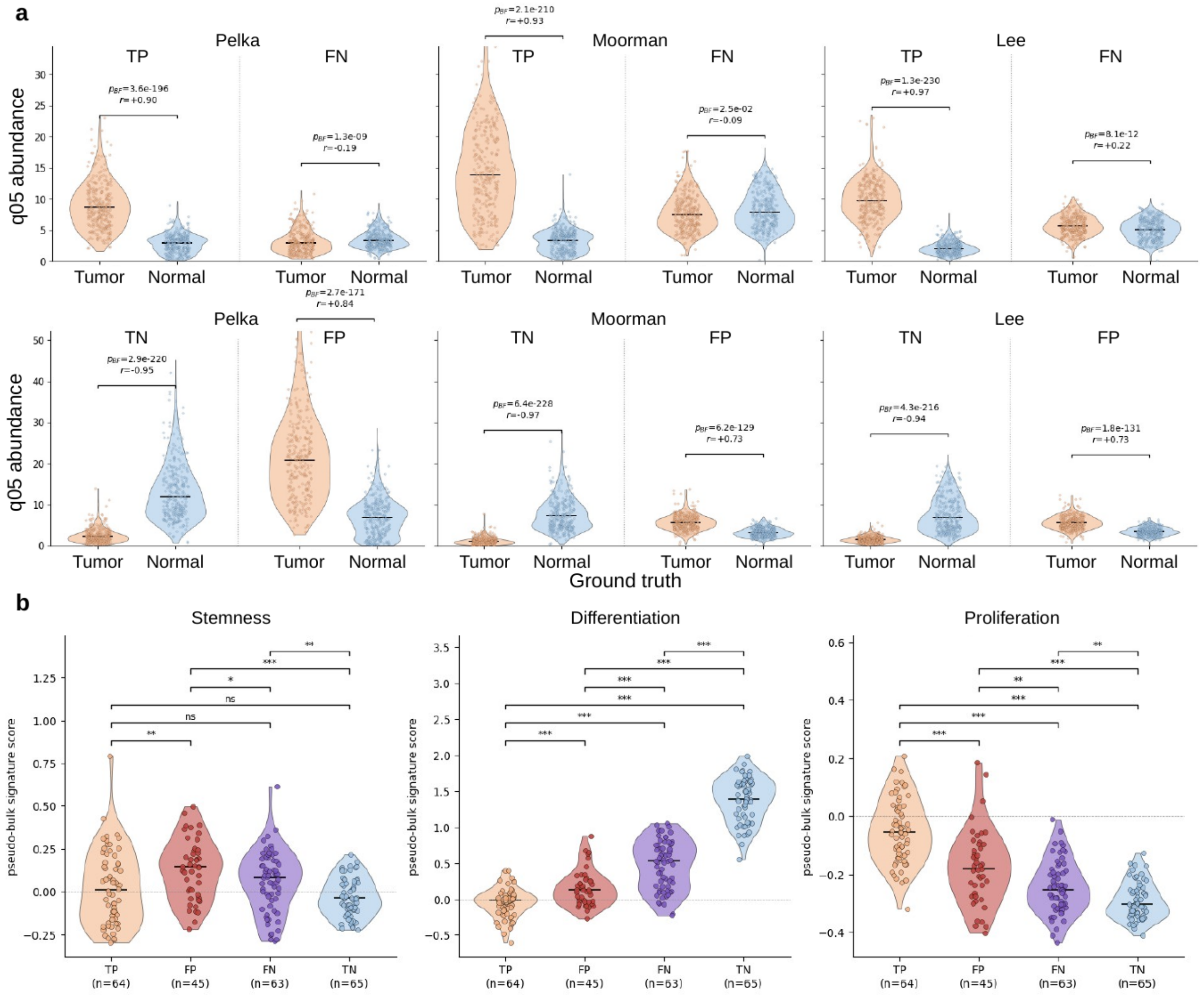


**Supplementary Fig. 7 | SPATNIC labels of tissue scRNA-seq cells agree with annotation and recover deconvolution-consistent spatial signal.** (a) Confusion matrices comparing original annotations with SPATNIC labels in three CRC scRNA-seq datasets: Pelka et al. 2021 (168,295 cells), Moorman et al. 2025 (34,955 cells), and Lee et al. 2020 (11,197 cells); counts with row-normalized percentages. (b) Cell2location deconvolution mapping of SPATNIC-defined TP, FP, FN, and TN cells onto matched spatial ground truth for each dataset. (c) q05 cell-abundance distributions of SPATNIC TP/FN and TN/FP populations stratified by ground-truth tumor/normal status across the three datasets; Spearman r and P values shown. (d) Pseudo-bulk stemness, differentiation, and proliferation signature scores across the TP, FP, FN, and TN groups; significance indicated.

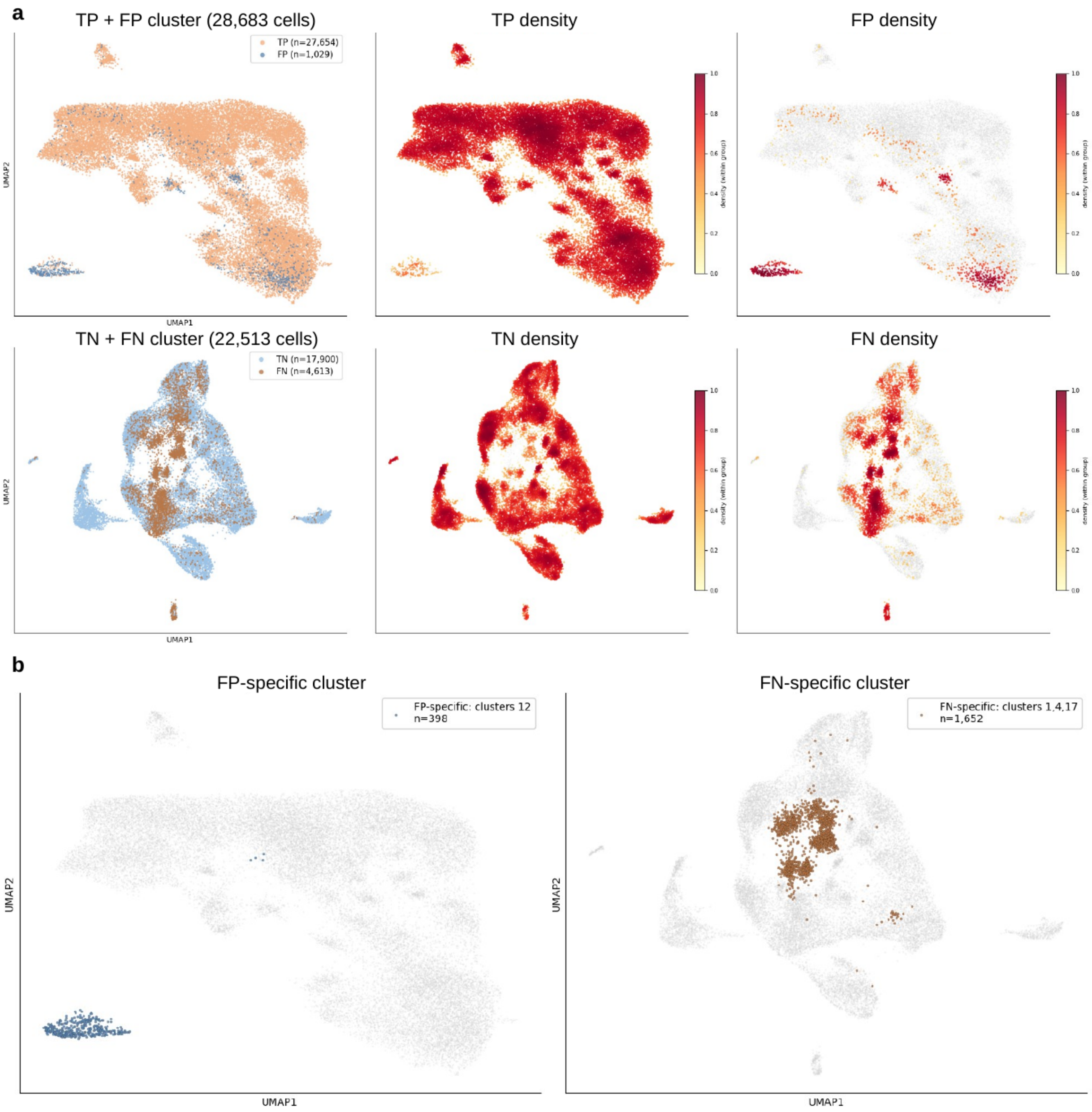


**Supplementary Fig. 8 | False-positive and false-negative calls localize to distinct transcriptional clusters.** (a) UMAP of combined TP+FP cells (28,683) and TN+FN cells (22,513), with accompanying density maps for TP, FP, TN, and FN populations; counts (n) per class indicated. (b) Identification of an FP-specific cluster (clusters 12; n = 398) and an FN-specific cluster (clusters 1, 4, 17; n = 1,652) highlighted on the UMAP.

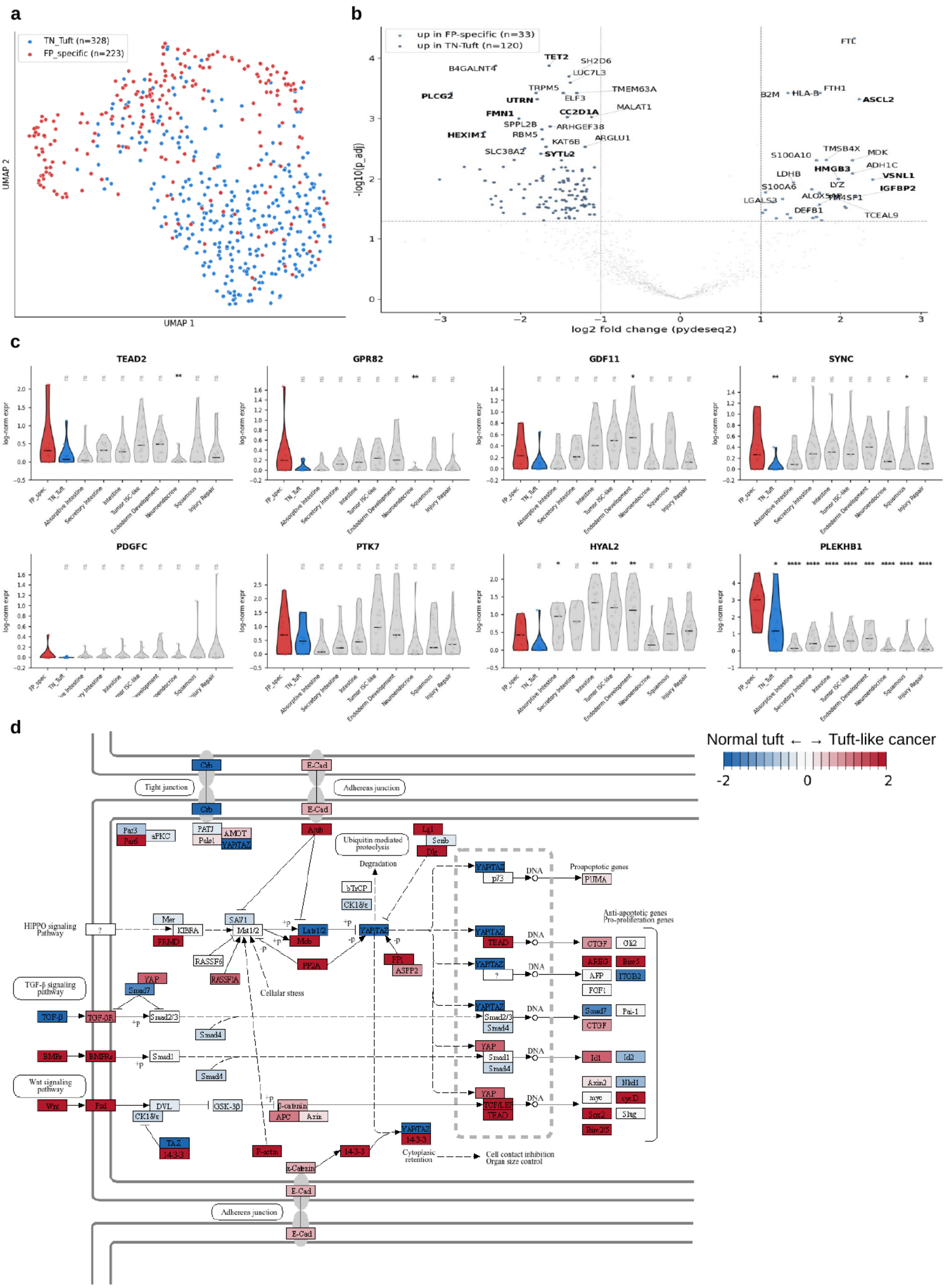


**Supplementary Fig. 9 | The FP-specific cluster is a *POU2F3*⁺/*ASCL2*⁺ tuft-like cancer population marked by Hippo/Wnt/TGF-β activity.** (a) UMAP of TN-tuft (n = 328) and FP-specific (n = 223) cells. (b) Differential expression between FP-specific and TN-tuft cells (pydeseq2); genes enriched in each group are labeled, including *ASCL2*, *HMGB3*, *IGFBP2* (tuft-like cancer) and tuft markers in the opposing direction. (c) Expression of selected discriminative genes (*TEAD2*, *GPR82*, *GDF11*, *SYNC*, *PDGFC*, *PTK7*, *HYAL2*, *PLEKHB1*) across the FP-specific and TN-tuft clusters relative to other epithelial subsets; significance indicated. (d) KEGG Hippo-signaling pathway diagram colored by the FP-specific (tuft-like cancer) versus TN-tuft (normal tuft) expression contrast, highlighting coordinated TGF-β, Wnt, and Hippo activation.

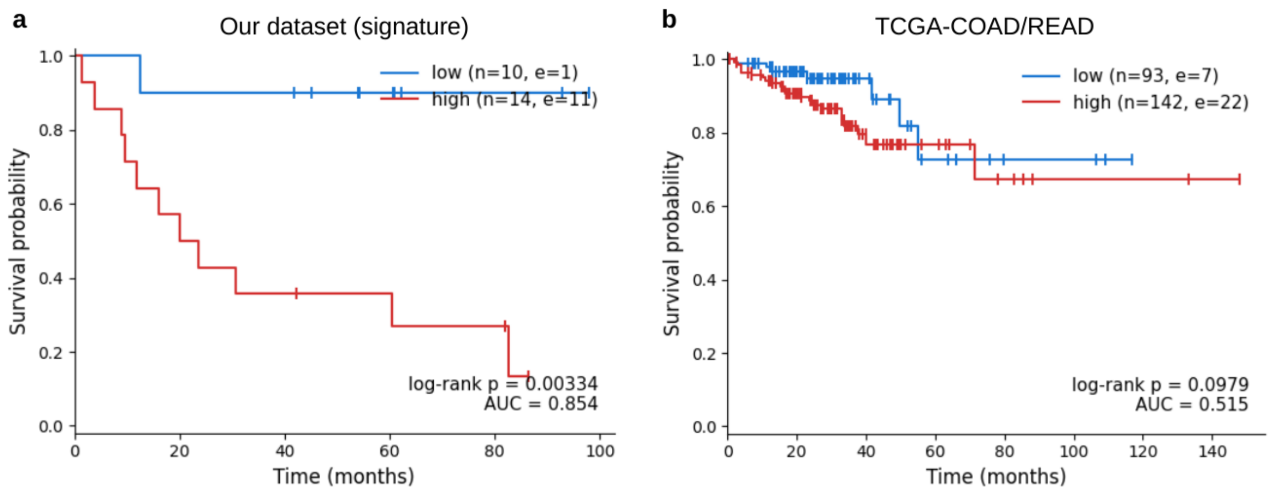


**Supplementary Fig. 10 | The tuft-like cancer signature stratifies relapse risk in an independent cohort.** (a) Kaplan–Meier survival by tuft-like cancer signature (high vs low) in the in-house dataset; log-rank P and signature AUC shown. (b) Corresponding analysis in TCGA-COAD/READ. Group sizes (n) and event counts (e) are indicated.

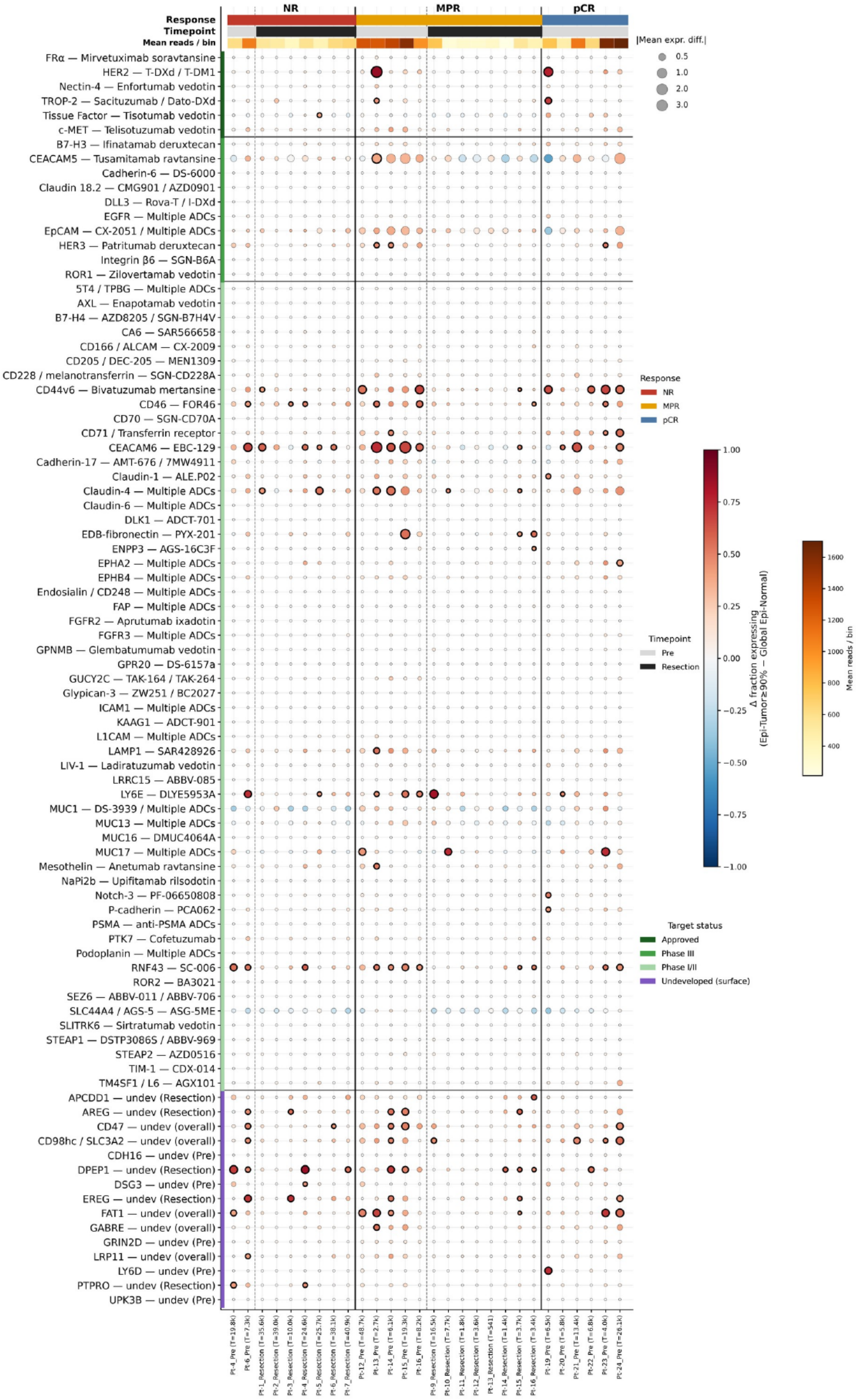


**Supplementary Fig. 11 | Full ADC-target dot plot across the LARC Visium HD cohort.** Differential expression of all 87 candidate antigens (Supplementary Table S8; 72 antigens targeted by clinically developed ADCs or antibody therapeutics and 15 undeveloped cell-surface candidates) between SPATNIC-defined Epi-Tumor and pooled (global) Epi-Normal compartments, stratified by treatment response (non-response, NR; major pathologic response, MPR; pathologic complete response, pCR) and timepoint (Pre, Resection). Samples were retained if their mean Epi-Tumor depth was at least 700 reads per bin (28 samples: 13 Pre, 15 Resection). Dot color, difference in the fraction of expressing bins (Epi-Tumor minus global Epi-Normal); dot size, absolute mean expression difference (log-normalized); bold outline, significant differential expression (Benjamini-Hochberg FDR < 0.05 and AUROC >= 0.7, Tumor > Normal). The top color strip shows the per-sample mean number of reads per 16-um bin. Rows are grouped by target development status (approved, phase III, phase I/II, and undeveloped surface candidate). The main figure (Fig. 5b) shows a curated subset of these antigens.
